# Genetic Evidence Indicates the Evolutionary Importance of the SARS-CoV-2 ORF9b protein

**DOI:** 10.64898/2026.02.23.707522

**Authors:** Ryan Hisner, Darren P Martin

**Affiliations:** Division of Computational Biology, Department of Integrative Biomedical Sciences, Institute of Infectious Diseases and Molecular Medicine, University of Cape Town, Cape Town, South Africa

## Abstract

All known betacoronaviruses possess an overlapping alternate-frame gene within the nucleocapsid gene. In SARS-CoV-2, the gene for this “internal protein” is ORF9b. The WHO Variants of Concern (VOC) Alpha, Delta, and Omicron all possess noncoding mutations that increase ORF9b expression. We show that with two exceptions, every major variant of the VOC era has had similar noncoding mutations to increase ORF9b expression and that these mutations are also frequently seen in long-branch, anachronistic, posited chronic-infection (PCI) sequences. Furthermore, we show that the amino acid substitution rate in ORF9b is higher than for any other SARS-CoV-2 gene, both in high-quality circulating sequences and in PCI sequences. This suggests that, as immunity to SARS-CoV-2 has grown in the population, increased ORF9b expression has conferred an evolutionary advantage, likely due to its ability to antagonize the antiviral type-I interferon response. We also show that PCI sequences are marked by distinct mutational patterns in ORF9b, some of which later became prominent in major variants. This evidence points to the importance of ORF9b for both viral transmission and within-host persistence.

## Introduction

One of the hallmarks of COVID-19, the disease caused by SARS-CoV-2, has been a delayed, dysregulated innate immune response marked by severely blunted interferon (IFN) activation. The IFN response is crucial to slowing viral replication and preventing severe disease, and numerous studies have documented a correlation between poor early IFN responses and severe disease outcomes^1–3^.

One mediator of dysregulated early IFN responses is the viral protein, ORF9b. ORF9b is expressed from a subgenomic RNA (sgRNA) contained within the first 100 codons of the nucleocapsid (N) gene but translated in a +1 reading frame relative to N. Experimental evidence indicates that ORF9b has two distinct folded states: a monomer (hereafter referred to as ORF9b-mon), composed mostly of alpha helices that binds the mitochondrial membrane protein TOM70, and a dimer (hereafter referred to as ORF9b-dim) consisting of beta sheets^4–7^ (refs). Whereas ORF9b-dim is likely involved in virion assembly^4, 8–9^ (refs), the monomer form is likely involved in dysregulating IFN responses^7, 10–11^ (refs). In SARS-CoV, ORF9b-dim associates with virions, and the ORF9b homolog in the JHM strain of mouse hepatitis virus (JHMV), called the I protein, is a structural protein essential for efficient viral assembly^8–9, 12^ (refs). Although all other known members of the Betacoronavirus subgenus to which SARS-CoV-2 belongs also encode ORF9b homologues in the +1 reading frame relative to their N genes, the structures, forms, interactions and function of these homologues remain largely unexplored.

ORF9b is not even annotated in the SARS-CoV-2 reference genome and ORF9b mutations are not listed on GISAID sequences or the global SARS-CoV-2 phylogenetic tree. Nevertheless ORF9b elicits a stronger host antibody response than most other SARS-CoV-2 proteins. Assessment of convalescent serum from people infected with pre-variant of concern (VOC) SARS-CoV-2 lineages found IgG and IgM antibodies to just four of the 18 proteins tested, with antibodies to ORF9b being third-most abundant, trailing only N and spike^13–14^. Further, ORF9b antibodies have also been detected in SARS-CoV-1 convalescent serum, and in the serum of calves infected with bovine coronavirus^15–16^.

Given the uncertainty surrounding the functions of SARS-CoV-2 accessory proteins, genetic and epidemiological evidence that has steadily accumulated since the start of the pandemic may prove vital to revealing the contributions of these proteins to SARS-CoV-2 fitness at different stages of the pandemic, illuminating the evolutionary dynamics of the genes encoding these proteins to highlight which of them deserve more intensive study, and guiding decisions on which therapeutic targets are likely to be most worth pursuing.

Here we present genetic evidence of the important, but under-appreciated, role of ORF9b evolution in the adaptation of SARS-CoV-2 to humans. We present evidence of multiple convergent ORF9b-associated genetic changes in dozens of the most successful circulating SARS-CoV-2 lineages and hundreds of individual highly divergent sequences likely originating in chronic infections of immunocompromised individuals. These signals of convergent evolution are some of the most remarkable seen during the SARS-CoV-2 pandemic and support the hypothesis that genetic changes to ORF9b and the genome sites that impact its expression levels were among the most crucial evolutionary adaptations enabling the emergence of almost all major circulating SARS-CoV-2 lineages since 2020.

## Methods

### Sequences datasets

We used an iterative approach to assemble a dataset of sequences from the GISAID database^17^ that had most likely arisen in a chronic infection context. An initial posited chronic-infection (PCI) sequence dataset was assembled manually over the course of over two years between April 2022 and January 2025. Candidate chronic-infection sequences were identified through (i) manual examination of individual sequences and sequence metadata on GISAID, (2) manually scanning 5000-sequence subsamples of the most up-to-date global SARS-CoV-2 phylogenetic tree for sequences separated from the rest of the phylogeny by unusually long branches, and (iii) manual examination of sequences on Nextclade^18^. After collecting several hundred such sequences, manual searches on GISAID and CovSpectrum^19^ for mutations found to be common in PCI sequences (present in >2% of the sequences) but rare in circulating lineages (present in <0.1% of sequences) were performed. ORF1a:K1795Q, for example, is present as a private mutation in 9.2% of the PCI sequences but <0.01% of all high-quality circulating sequences (excluding the Gamma VOC lineage and lineage BS.1) as of 2025-01-12 (Supplementary Table S1). Approximately 1200 additional sequences carrying mutations that differentiated the PCI sequences from circulating sequences were added to a preliminary expanded posited chronic-infection (EPCI) dataset together with the original set of PCI sequences.

Nextclade CLI was used to create ndjson files of the EPCI sequence collection. Custom written Julia code (https://github.com/ryhisner/ORF9b_relevant_code) was then used to identify and count the private mutations for each sequence (with private mutations being defined by Nextclade as “the mutations between the query sequence and the sequence corresponding to the nearest neighbor (parent) on the tree.”). Closely related sequences very likely collected from a single individual were identified by a combination of geographical location (same country), shared Pango lineage designations, and six or more shared private mutations. Further, sequences carrying >8 reversion mutations were excluded because these would have likely been either low-quality sequences or recombinants. The phylogenetic placement of the EPCI sequences in the global SARS-CoV-2 tree was examined using USHER^20^ and clusters of EPCI sequences within this tree were manually examined using Nextclade web^18^ to determine whether sequences in these clusters all had a recent common origin or had independently evolved but were spuriously clustered within the global SARS-CoV-2 tree due to presumed long branch attraction^21^.

For groups of related sequences (typically from the same individual), the sequence with the most private amino acid mutations was chosen as the single representative for each group. This yielded an interim EPCI dataset comprising 3483 sequences each from a different potential chronic infection (https://github.com/ryhisner/EPCI_dataset).

### Identification of convergent mutations

Custom written Julia code (https://github.com/ryhisner/ORF9b_relevant_code) was used to analyze a Nextclade ndjson file containing information from all sequences within the EPCI dataset. Using the mutations identified by Nextclade as private, the most common mutations in these sequences were identified. To identify mutations that are distinctive to chronic-infection sequences (i.e. those mutations only infrequently seen in circulating lineages but which are relatively common in the EPCI sequences), a Nextclade ndjson file for all SARS-CoV-2 sequences on GISAID was created. Using a strict quality control filter (Nextclade overall qc score ≤ 5, zero unrecognized frameshifts, ≤5 private amino acid substitutions), private mutations in circulating lineages were counted. Artifactual reversions are extremely common in SARS-CoV-2 sequences due to sequence assembly pipelines commonly defaulting to the reference genome in regions of the genome with high degrees of sequencing failure, while genuine reversions are exceedingly rare. Reversions were therefore excluded from the analysis of circulating lineages.

For each mutation, a ratio of the number of times it appeared as a private mutation in the EPCI sequences to the number of times it appeared as a private mutation in circulating lineages was calculated (chronic-to-circulating mutation ratio, CCMR). This ratio was normalized by multiplying it by the ratio of the total number of amino acid mutations in high quality circulating sequences (HQCS) to the total number of amino acid mutations in EPCI sequences. Mutations with the highest ratios by this measure and which occur at least three times independently were identified as being “convergent” in EPCI sequences.

Using the ratio described above multiplied by the number of times a mutation independently occurred among sequences in the list, a “convergence score” was calculated for each mutation. Mutations that appeared at least twice and which either had convergence scores above 125 or CCMR > 10 were added to a “convergent mutations” list. This list was then used to identify additional sequences deemed likely to have originated in chronic infections which were not already within the EPCI dataset. Specifically, ndjson files containing subsets of one million SARS-CoV-2 GISAID accession numbers (EPI_ISL_8000001-EPI_ISL_9000000, for example) were then scanned for sequences that contained (i) at least four private mutations from the “convergent mutations” list (Supplementary table S1), (ii) at least eight private mutations overall, (iii) not more than two reversions, and (iv) fewer than 120 private mutations. The vast majority of the additional tens of thousands of potential chronic-infection derived sequences identified in this manner were extremely low-quality sequences, often containing mutations from multiple clades. These sequences were manually removed, and the remaining sequences were then assessed individually. Using USHER, sequences that were placed on long branches (≥6 non-synonymous substitutions, deletions, or insertions) and which were collected more than two months later than the most closely related sequences on the global SARS-CoV-2 phylogenetic tree were then added to the EPCI dataset. For borderline cases, sequences for which ≥80% of nucleotide substitutions in coding regions were non-synonymous were included and those with lower percentages excluded. This yielded the final EPCI dataset comprising 3297 sequences each from a different potential chronic infection (https://github.com/ryhisner/EPCI_dataset). We must stress that this EPCI dataset is intended as an enrichment of chronic-infection derived SARS-CoV-2 and does not represent a set of sequences that are provably derived from long-term chronic infections.

### Comparisons of mutation dynamics in different SARS-CoV-2 genes and gene regions

To calculate a “raw mutational density” (RMD) score (i.e. the rate at which mutations occur) for each gene in the EPCI sequences (adjusted for gene length), the total number of private amino acid substitutions in each gene was counted and divided by the number of amino acid residues contained in the corresponding gene.

A “chronic-to-circulating substitution ratio” (CCSR) was calculated by dividing the RMD score of each gene in the EPCI dataset by the RMD for each gene in the HQCS dataset. This ratio was then adjusted by multiplying it by the total number of private mutations in the HQCS dataset divided by the total number of private mutations in the EPCI dataset.

To measure the mutational density in sub-gene regions, the same method described above was used but, instead of focusing on entire genes, sliding 11-amino acid-long sequence windows spanning each gene at 1-AA intervals were individually examined. The raw mutation counts and CCSR can be found in Supplementary table S2.

### Structural Data

ORF9b structures were visualised using ChimeraX^22^. ORF9b dimer structures (PDB 6Z4U, PDB 7YE7, and PDB 2CME) and OR9b-TOM70 structures (PDB 7DHG and PDB 7KDT) were obtained from the Protein Data Bank.

### Results and Discussion

Among the known betacoronaviruses, internal, overlapping open reading frames within the N gene are a conserved feature, yet the functions, forms and structures of the proteins these ORFs encode are mostly unknown. Structures only exist for the SARS-CoV and SARS-CoV-2 ORF9b proteins, both of which exist in structurally disparate monomer (ORF9b-mon) and dimer (ORF9b-dim) forms. Despite having very little sequence homology, the predicted structures of ORF9b homologs in MHV-JHM and MERS-CoV closely resemble those of the betacoronavirus ORF9b-mon proteins^1, 8^. As in SARS-CoV and SARS-CoV-2, the MHV-JHM and MERS-CoV ORF9b homologs antagonize IFN induction^1, 8, 23^. In MERS-CoV this protein is expressed in multiple different isoforms, with the translation of each isoform initiated from a different start codon^23^.

Overlapping genes generally have lower mutation rates than non-overlapping genes^24^, yet ORF9b has been one of the most evolutionarily dynamic genes in SARS-CoV-2, both during the diversification of circulating lineages, and during intra-host evolution in immunocompromised individuals. This evolution has been marked by striking convergence, across multiple independent viral lineages, converging upon a particular mutational pattern that has not been seen in any previous betacoronavirus genomes.

### Mutations that increased the production of N/ORF9b subgenomic RNAs were likely strongly favoured by natural selection during the VOC era

As with other coronaviruses, SARS-CoV-2 produces a variety of different mRNAs for structural and accessory gene expression via an unusual template switching mechanism. As the RNA-dependent RNA polymerase (RdRp) traverses the positive-sense genome, from the 3’ to 5’ end, it encounters a number of transcription regulation sequences (TRSs) that match a “leader” TRS, called the TRS-L, in the 5’ UTR of the genome^25^. Each of these “body” TRSs, which are collectively called TRS-Bs, precedes a different open reading frame (ORF) from which one or more subgenomic mRNAs (sgmRNAs) are produced. During synthesis of complementary RNA strands, the RdRp pauses at TRS-B sequences. In a certain proportion of instances during these pauses, determined to some extent by the degree of similarity between sequences surrounding the TRS-L and TRS-B sequences, the RdRp will either continue through the TRS-B and onwards, or jump from the TRS-B to the TRS-L of the genomic RNA molecule to yield a negative-sense subgenomic RNA (sgRNA-) molecule. Positive sense sgmRNAs that are then synthesised from the various sgRNA-molecules will include a variety of different genes attached directly to the same leader sequence. These capped and 2’-O-methylated sgmRNAs constitute a heterogeneous population of mRNA molecules that are collectively primed for translation into the entire spectrum of coronavirus proteins^26^.

The length and degree of TRS-L homology varies for different TRS-Bs, with more extensive homology generally leading to more frequent template switching and higher rates of sgRNA production. The “core” or minimal TRS-L for SARS-CoV-2 is ACGAAC, but the TRS-Bs for the two most highly expressed sgmRNAs, those expressing the membrane (M) and nucleocapsid (N)/ORF9b proteins, respectively, match eleven (TCTAAACGAACT) and twelve (TCTAAACGAAC) nucleotides from the TRS-L region. Conversely, the TRS-B for the envelope (E) protein, which is expressed at very low levels, features only the minimal ACGAAC TRS motif.

The distance of each gene from the 3’ end of the genome also affects the relative production levels of different sgmRNAs, with genes closer to the 3’ end tending to be more abundantly produced than those closer to the 5’end of the genome.

In SARS-CoV-2 the TRS-B associated with the N/ORF9b genes is particularly interesting in that it is a “double” TRS-B. Specifically, the N/ORF9b-associated TRS-B consists of a slightly suboptimal, secondary 3’ TRS overlapping an optimal, primary 5’ TRS core (Figure 1). This, together with its position as the 3’-most TRS-B in the genome and possibly other factors (such as its unique position on the 3’ side of a stem-loop, where every other TRS-B appears on the 5’ side of a stem-loop)^27^, combine to make the N/ORF9b sgmRNA by far the most abundant of all those produced by SARS-CoV-2^28–30^. The reasons for this likely relate to the importance of both N and ORF9b expression during the early phases of infection.

**Figure 1:**
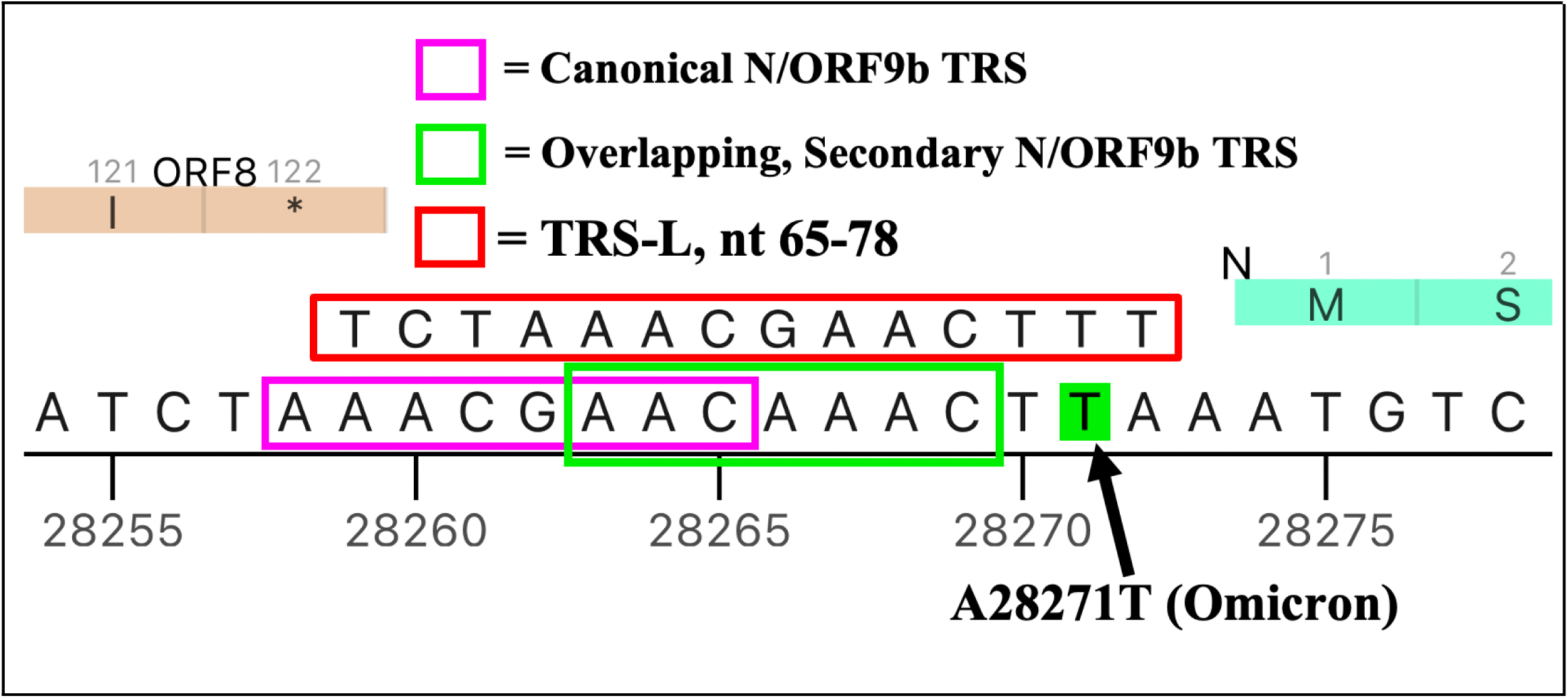
Overlapping transcription regulation sequence (TRS) for the nucleocapsid/ORF9b sgmRNA. In addition to dramatically decreasing the favorability of the N Kozak sequence, A28271T also creates additional extended homology with the TRS-L

The evolutionary benefit of maintaining optimal N/ORF9b sgmRNA production in SARS-CoV-2 is likely reflected by the fact that the primary N/ORF9b associated TRS-B and its immediate flanking nucleotides have remained highly conserved since the start of the pandemic. The only mutation in this region in any major SARS-CoV-2 lineage was a remarkable “AACA” insertion in the Gamma VOC lineage after nucleotide 28262. This insertion created a “triple” TRS-B, adding yet another overlapping, slightly suboptimal TRS (Figure 2): a modification that would be expected to increase N/ORF9b sgmRNA production.

**Figure 2:**
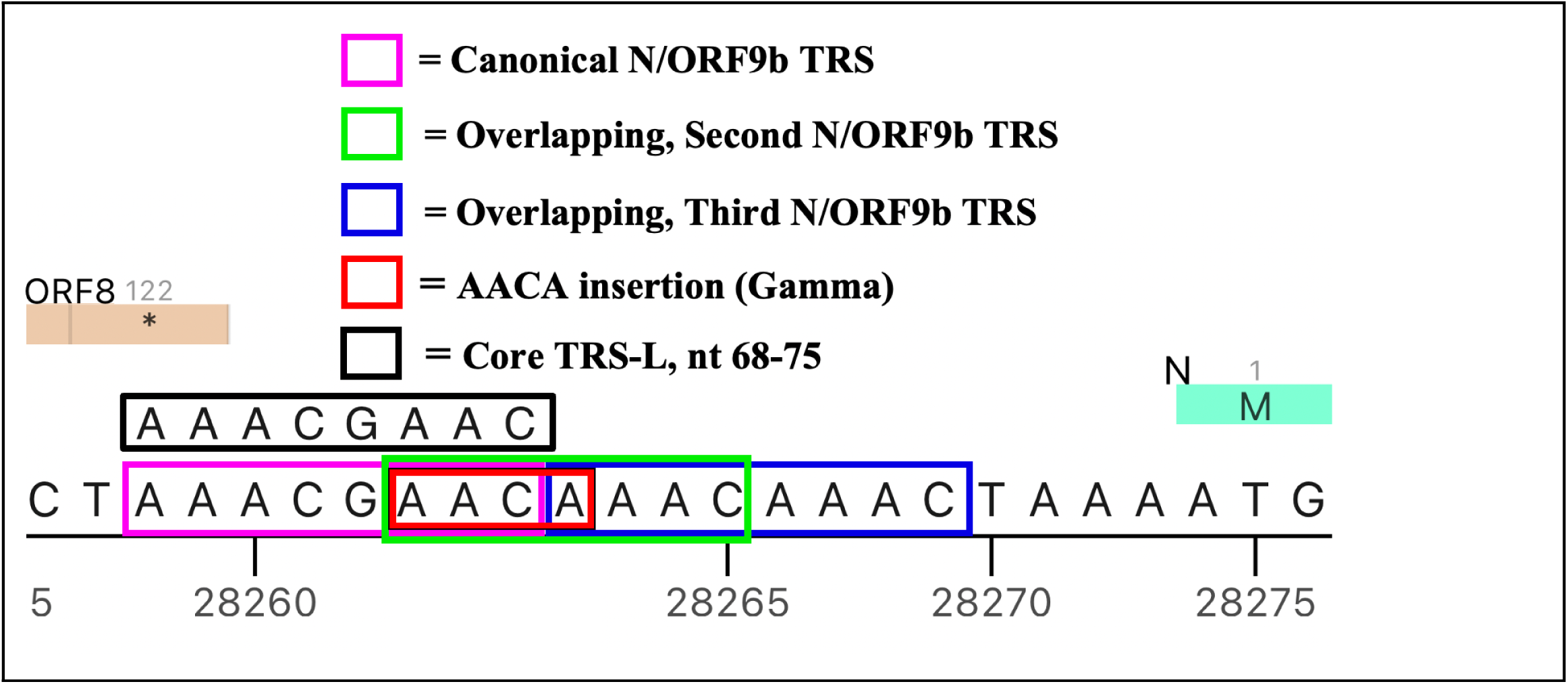
The AACA insertion in the Gamma VOC creates a third overlapping N/ORF9b TRS.

Though pre-VOC SARS-CoV-2 lineages were not known to possess a functionally significant ORF9b-specific TRS-B (i.e. a TRS-B that would yield a sgmRNA from which only ORF9b could be expressed), these early lineages possessed a vaguely TRS-like motif that overlaps the ORF9b start codon (Figure 1). The Alpha VOC lineage characteristically possesses a peculiar three-nucleotide mutation (resulting in a N:D3L amino acid change), which increases the similarity of this TRS-like motif to the TRS-L so that eight nucleotides perfectly match the 5’ region of the TRS-L (TCT**<u>CTA</u>**AA) while 11/13 nucleotides match the full, 5’-extended TRS-L (TCT**<u>CTA</u>**AA*T*G*G*AC—substitutions causing the N:D3L amino acid change underlined bold, non-TRS-L-matching nucleotides in italics). Accordingly, the N:D3L mutation results in a 6- to 80-fold increase in the production of ORF9b-specific sgmRNA^31, 89^. However, the vanishingly small levels of the ORF9b-specific sgmRNA produced by pre-Alpha isolates mean that this increased ORF9b-specific sgmRNA production still only accounts for 0.5% of all the sgmRNAs produced by Alpha isolates. Further, the increased ORF9b-specific sgmRNA production that occurs in N:D3L mutants does not, on its own, result in increased levels of ORF9b protein expression^32–33, 89^.

### Mutations favoring ORF9b expression over that of N were also likely strongly favoured by natural selection during the VOC era

In addition to evidence of increased ORF9b expression being strongly favoured by TRS mutations during the first three years of the pandemic, there is also evidence that selection has strongly favoured mutations within the N gene Kozak sequence that specifically enhanced ORF9b expression. The Kozak sequence of a gene consists of the nucleotides surrounding its start codon. The identity of the nucleotides at each position can affect the rate at which ribosomes recognize the start codon and begin translation. In particular, the -3 position (the third nucleotide on the 5’ side of the start codon) has the largest effect on translation efficiency in humans, with purines (A/G) creating a much more favorable context for translation than pyrimidines (C/T).

ORF9b is produced from the same sgmRNA that produces N through a process referred to as leaky scanning, where a ribosome bypasses the first start codon in an mRNA. The frequency of leaky scanning is largely determined by the favorability of the Kozak sequence of the first start codon^34–36^.

In the SARS-CoV-2 Wuhan-Hu-1 reference genome, the Kozak sequence for N is favorable (though not perfect), featuring the most favorable nucleotide (A) at the most impactful -3 position but lacking a G in the +4 position, the second most important Kozak site^34^ (Figure 2; ref).

In addition to carrying an N:D3L mutation that enhances ORF9b-specific sgmRNA production, the Alpha VOC lineage also carries a mutation in its N Kozak sequence: a single nucleotide deletion at genome position 28271 (ΔA28271/A28271del). This deletion changes the crucial -3 Kozak-sequence residue from the most favorable (A) to the least favorable nucleotide (T), the largest possible decrease in Kozak favorability with a single nucleotide change. Accordingly, the quantity of ORF9b protein produced by Alpha exceeds that of pre-VOC variants by 6 to 7-fold, both *in vitro* and in infected Calu-3 human lung cells^31, 33, 38, 89^. Surprisingly, despite its drastically reduced Kozak favorability, Alpha isolates still produce more than twice as much N protein as pre-VOC lineages^31, 38^.

Remarkably, the Delta VOC lineage, which became dominant worldwide between January and May 2021 when it displaced all previous variants, had this same ΔA28271 deletion, resulting in an increase in ORF9b protein expression similar to that of Alpha^89^..

Later, the various Omicron lineages that emerged in late 2021 and early 2022 (BA.1, BA.2, BA.4, and BA.5) all shared a A28271T mutation which causes an even larger decrease in Kozak favorability than ΔA28271^39^ (due to a modest compensatory increase in Kozak favorability at -4 with ΔA28271), together with a slight increase in extended TRS-L homology for the “secondary” N/ORF9b TRS-B (Fig 1).

In stark contrast to convergent spike mutations seen throughout the pandemic, this remarkable genetic convergence in three major lineages that were either globally dominant (Omicron, Delta) or dominant in a large number of countries (Alpha) has, with a few outstanding exceptions, been largely overlooked.

Still more overlooked is the striking presence of similar, likely ORF9b-upregulating, N-Kozak mutations in other VOC lineages, variants of interest (VOI), and numerous other significant lineages. Apart from the Beta and Gamma VOC lineages, every other VOC/VOI lineage has possessed a mutation predicted to decrease the favorability of the N gene Kozac sequence, and which would therefore very likely have increased leaky scanning and ORF9b protein expression (Table 1)^34–36^.

**Table 1.**
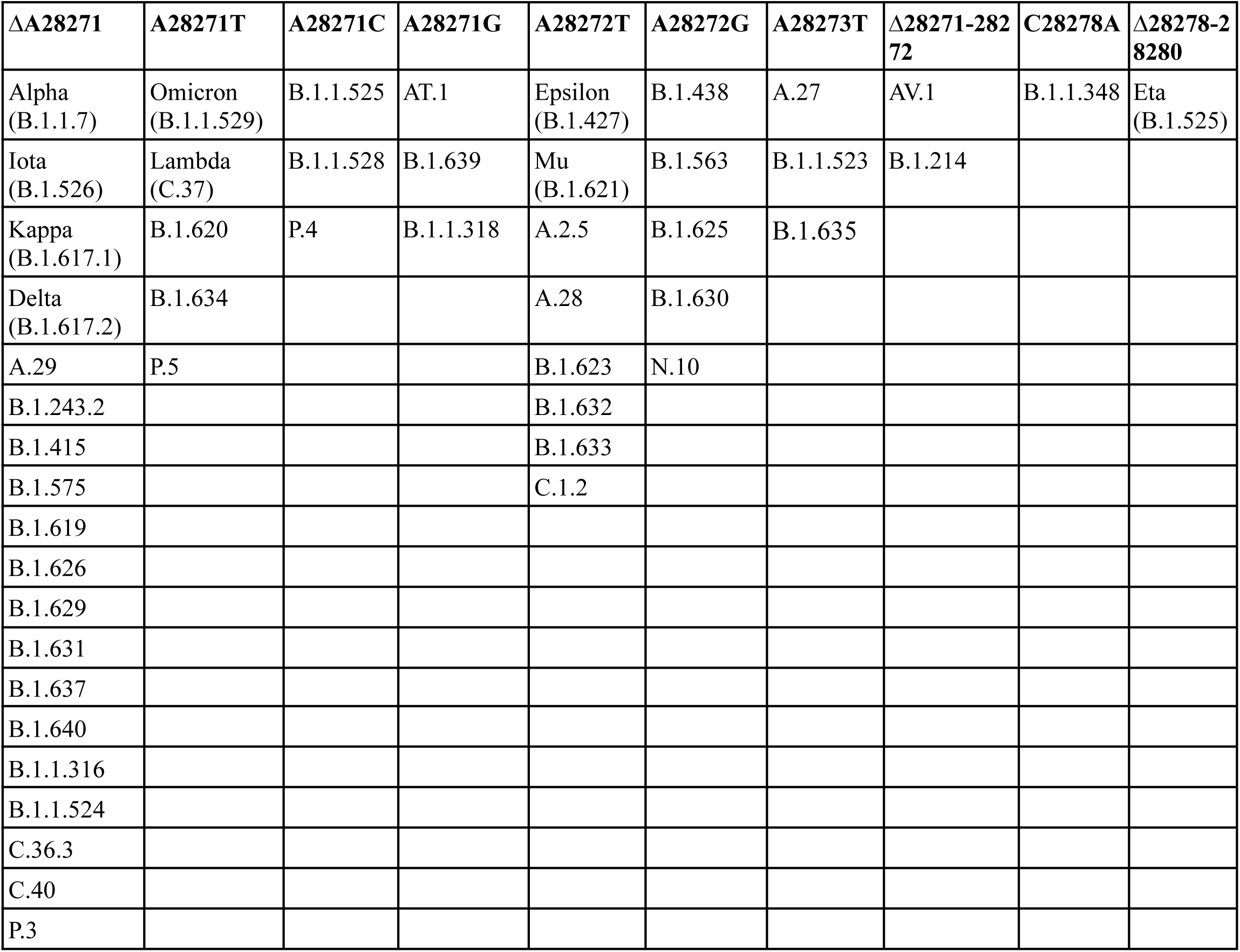
N-Kozak mutations characterizing major SARS-CoV-2 lineages during the VOC era of the pandemic.

In addition to Alpha and Delta, Iota (B.1.526), Kappa (B.1.617.1), and B.1.640 (the only pre-Omicron variant to compete well with Delta) had ΔA28271. Lambda (C.37) and B.1.620 had a convergent A28271T mutation with all Omicron variants while Epsilon (B.1.427/429), Mu (B.1.621), and C.1.2 all possessed A28272T. Eta (B.1.525) had Δ28278-28280, which causes a modest decrease in the favorability of the N-gene Kozak sequence^37, 40^ (refs). A large number of all other relatively successful and/or genetically significant variants had similar mutations expected to increase ORF9b expression (Table 1).

Apart from being among the most common mutations defining the VOC and VOI lineages, the importance during ongoing SARS-CoV-2 evolution of mutations that decrease the favorability of the N-Kozak sequence is also strongly supported by overall mutational patterns seen throughout the pandemic. Starting from the Wuhan-Hu-1 reference genome, there are roughly equal numbers of possible mutations between positions -4 and +6 spanning the N start codon that would be expected to increase (eleven mutations) and decrease (ten mutations) N-Kozak favorability. Yet, no Pango-designated lineage has ever had a mutation increasing N Kozak favorability. Furthermore, there have been no detected reversions at T28271 to the A residue found in Wuhan-Hu-1. T28271C, which causes a very slight increase in N Kozak favorability, has been seen in just 0.22% of sequences since January 1, 2022 (over 75% of which are BF.11 and its sublineages). Furthermore, 40.3% (104/258) of our EPCI sequences from clades that lack N-Kozak mutations possess a private N-kozak mutation predicted to increase leaky scanning and therefore enhance ORF9b expression.

Collectively these various N-Kozak mutations (Table 1), all of which are likely to enhance ORF9b expression, have involved dozens of lineages, including nearly all the most successful variants in the VOC era (the only major exceptions being Beta and Gamma) and are arguably the most extraordinary instance of SARS-CoV-2 convergent evolution that we have yet seen. These mutations certainly deserve mention alongside better known examples of SARS-CoV-2 evolutionary convergence, such as S:Δ69-70, S:E484K, S:N501Y, NSP6_Δ106-108 (ORF1a:Δ3675-3677), and N:R203K-G204R^41–43^.

Crucially, the sequence evidence that increased ORF9b expression relative to early pre-VOC SARS-CoV-2 lineages has likely provided more recent lineages with a decisive selective advantage is supported by direct measurements of absolute protein abundance during infection. While such measurements are difficult, there are several indications both that ORF9b is one of the most highly expressed proteins (only surpassed by M and N in the ancestral D614G lineage)^28^, and that its expression levels have increased substantially in VOC lineages relative to pre-VOC lineages. In fact, in VOC lineages ORF9b protein production has increased more relative to pre-VOC lineages than any other SARS-CoV-2 protein^33, 38, 89^. This is true even for Beta and Gamma which lack the Kozak-sequence mutations found in all other VOC lineages^89^.

### The N-Kozak sequences of SARS-CoV-2 have become uniquely suboptimal relative to those of other known betacoronaviruses

All other betacoronavirus ORF9b homologs are also thought to be translated through leaky scanning. The N-Kozak sequences are highly conserved among betacoronaviruses. Every known non-SARS-CoV-2 betacoronavirus genome is characterized by the good-but-not-perfect Kozac context frequently found in genes that allow a small degree of leaky scanning^34^. Without exception, they possess either the ideal A at position -3 or the ideal G at position +4, but not both. All but three currently known non-SARS-CoV-2 betacoronavirus genomes (two hibecovirus genomes, NC_025217.1, KF636752.1, and one unclassified betacoronavirus genome, OK017908.1) have either A or G at the -3 position, both of which are far more favorable than C or T. Multiple mutagenesis studies have now shown that substitution of T or C for A at the -3 position leads to increased production of both ORF9b^32–33, 89^ (refs) or ORF9b homologues^44^. Pre-VOC SARS-CoV-2 sequences had N Kozak sequences closely resembling those of other betacoronaviruses, with the optimal A at -3 but suboptimal nucleotides in other positions.

With the rise of Alpha and other early VOC and VOI lineages, suboptimal N-Kozak sequences first became prominent, and with the successive rise to global dominance of the Delta and Omicron VOC lineages, the SARS-CoV-2 N-Kozak sequence became uniquely suboptimal relative to those other known betacoronaviruses, a trend consistent with the exertion of intense selection pressures in humans favouring leaky scanning and greater ORF9b expression.

### The likely role of ORF9b-mon in IFN antagonism

A plausible reason that increased ORF9b expression in VOC lineages relative to pre-VOC lineages might be selectively advantageous in humans was presciently identified months before the emergence of the VOC lineages by Gordon et al^7^ (Gordon, 2020) and Jiang et al^11^ (Jiang, 2020). They identified the interaction of ORF9b-mon with the host mitochondrial membrane protein, TOM70. This interaction was associated with potent antagonism of the type-I IFN response induced by RIG-I-like receptors (RLRs).

RIG-I is the primary RLR that is active in SARS-CoV-2 infections. It detects 5’ triphosphates and double-stranded RNA, a common RNA-virus replication intermediate, leading to activation of the RIG-I CARD domains and oligomerization of RIG along the RNA molecule^45^. Oligomerized RIG-I in turn binds the CARD domains of the intracellular membrane protein, MAVS, causing it to undergo prion-like aggregation^46^. MAVS then further associates with TOM70, which recruits the chaperone protein HSP90. HSP90 ferries TBK1 and IRF3 to the mitochondrial surface, where IRF3 is phosphorylated, prompting it to migrate to the nucleus, where it activates genes encoding type-I IFN.^45–46^ Type-I IFN subsequently interacts with IFN receptors locally and in surrounding cells, setting off the JAK-STAT signaling cascade that leads to the expression of hundreds of IFN-stimulated genes (ISGs) which, through myriad mechanisms, induce an antiviral state^90^.

Antagonism of this IFN-response pathway by ORF9b-mon strongly suggests that the selective advantage of increased ORF9b expression in SARS-CoV-2 has been, in part at least, related to the suppression of type-I IFN responses. The enhanced suppression of these responses afforded by increased ORF9b expression may therefore have been one of the most important themes of SARS-CoV-2 evolution preceding the emergence of the Omicron lineages. It must be stressed, however, that ORF9b likely has additional, less well characterised, functions related to virion assembly; the shifting favorability during the pandemic of increased ORF9b expression in the context of these other functions is more difficult to discern.

### The functions, if not also the forms, of ORF9b-dim are likely conserved in diverse coronavirus ORF9b homologues

Given that ORF9b overlaps N and is therefore expected to be more evolutionarily constrained than non-overlapping genes, it is very interesting that it appears to have evolved the capacity to fold into two distinct forms: the beta sheet dominated ORF9b-dim form that appears to be involved in virion assembly and the alpha helix-based ORF9b-mon form that appears to be involved in IFN antagonism. There are only approximately 100 known fold-switching proteins, and fewer than 20 of those undergo an alpha-helix-to-beta-sheet transformation^47^. From both a structural and evolutionary perspective, ORF9b is therefore among the most unusual of all known proteins.

In both SARS-CoV and SARS-CoV-2, the well-ordered regions of each ORF9b-dim protomer take the form of beta sheets wrapped tightly, in handshake fashion, around the other protomer. Perhaps importantly, both contain a lipid-binding tunnel that holds a 19-21-carbon lipid that projects prominently outward from both sides. Furthermore, the planar surface nearest the lipid-binding cavity has a positive surface charge. It has therefore been proposed that the projecting ends of the encased lipid and the positively charged interface both mediate interactions with cellular membranes^4–5^.

Whether or not ORF9b homologs also form dimers in other betacoronaviruses is unknown, but there are several hints that they might do so. The MHV I protein (an ORF9b homologue) can be observed as a “doublet” using SDS-PAGE, has been detected as a structural component of MHV virions^12^, and impairs virus assembly when deleted^8^.

In SARS-CoV-2 it is likely that the C-terminal domain of ORF9b is crucial for dimer formation, interactions with membranes, and whatever precise function ORF9b has in virus-assembly. Deletion of seven C-terminal ORF9b amino acids abolishes dimer formation^48^. The ORF9b homologues of MERS-CoV, bovine coronavirus and MHV all localize in the ER, golgi and/or ER-golgi intermediate compartments: the same locations as all other viral structural proteins (S, E, and M) that are involved in virion assembly^8, 12, 23^. The involvement of the C-terminal domains of ORF9b and its homologues in binding the membranes of these compartments is supported by the fact that, with MERS-CoV, deleting the 25 C-terminal domain amino acids of its ORF9b homologue (called ORF8b) abolishes its localization to the ER-Golgi intermediate compartment^23^. All of these observations suggest that these various ORF9b homologues recapitulate the characteristics of the SARS-CoV and SARS-CoV-2 ORF9b-dim, and might therefore also do so in the form of dimers.

ORF9b has a YPX_n_L motif (where X_n_ is any amino acid, the number of which may vary), which is known to bind the human ESCRT protein ALIX^49^. ESCRT proteins are used by many enveloped viruses for viral trafficking and egress, including HIV^52–53^ (refs), hepatitis C virus^54^, hepatitis A virus^55^, and others, and it has recently been shown that numerous betacoronaviruses, including SARS-CoV-2, utilize cellular ESCRT machinery for assembly and egress^50^. This motif is highly conserved both among circulating SARS-CoV-2 sequences (<2000 private mutations in ∼10 million HQCS, see Table S3) and among sequences in our EPCI dataset, in which only two sequences have a mutation erasing the YPX_n_L motif—one of which occurs after a stop codon, while the other is part of an AATAATAAT repeat, likely a result of RdRp stuttering, which usually occur in A/T-rich nucleotide stretches^51^.

The ORF9b of human coronavirus OC43, despite sharing almost no sequence homology with the SARS-CoV-2 ORF9b, has a nearly identical YPX_n_L motif, which is also conserved in bovine coronavirus and many other embecoviruses. Furthermore, the ORF9b protein of human coronavirus HKU1, which has little sequence homology with OC43 ORF9b, possesses one or two YPLL motifs. When the consensus ORF9b amino acid sequences for OC43, HKU1, and SARS-CoV-2 are randomly rearranged 100,000 times, the average number of YPXL and YPXXL motifs expected by chance alone is under 0.06 per sequence.

The preservation of the YPX_n_L motif in the ORF9b of the four major betacoronaviruses that infect humans (including SARS-CoV), when considered alongside evidence that ORF9b is both incorporated into virions in SARS-CoV, and essential for efficient assembly in JHMV, suggests that the SARS-CoV-2 ORF9b-dim could be involved in viral assembly, trafficking, and egress.

In many ways, the expression dynamics and roles of ORF9b during the SARS-CoV-2 infection cycle resemble those of the N protein. Both are very likely preferentially expressed early in infection, during which they are directed to intracellular membranes (mitochondria for ORF9b-mon, and double-membraned viral replication organelles for N)^56–60^ where they powerfully antagonize innate immune reactions (ORF9b-mon by blocking the IFN signaling cascade, N by interfering with stress granule formation)^56–57, 61^.

Later in infection, N migrates to the endoplasmic reticulum Golgi-intermediate compartment, the primary site of viral assembly, as might ORF9b-dim by analogy with its presumed homologues in other betacoronaviruses. It should be stressed, however, that it is presently unknown where ORF9b-dim localises during SARS-CoV-2 infections. Furthermore, the shift in behavior between early and late infection periods for both N and ORF9b is likely governed by their phosphorylation statuses^7, 62–65^.

### Why ORF9b mutations arising during chronic infections might illuminate the likely trajectories of future ORF9b evolution

A hallmark of SARS-CoV-2 evolution has been the emergence of highly divergent, anachronous variants, i.e. ones not descended from contemporary circulating variants. Abundant evidence indicates that in many, if not almost all, cases these variants evolve within chronically infected individuals^56–77^. These divergent variants have typically been marked by a large number of spike mutations, but ORF9b mutations have also featured prominently. Among the ORF9b mutations, those that convergently evolve in multiple independent lineages are particularly interesting from the perspective of identifying amino acid changes that have made the virus better at infecting humans. Further, of these likely adaptive convergent mutations, those that differentiate circulating lineages from lineages sampled from chronic infections are particularly interesting from the perspective of understanding differences in the selective processes at play during intra-host persistence and inter-host transmission. Specifically some selective pressures on ORF9b in circulating viruses may be relaxed during chronic infections whereas there may be specific selection pressures on ORF9b during chronic infections that are different and/or stronger than those acting on the gene in actively circulating viruses. Detectable differences in patterns of adaptive mutations in chronic and circulating viruses might provide some insights into the features and functions of ORF9b that differ between persistent and chronic infections.

We therefore assembled a dataset of SARS-CoV-2 of 3297 whole genome sequences that were likely sampled from long-term chronic infections: a dataset referred to as our EPCI dataset. Many mutations that have become predominant in circulating SARS-CoV-2 sequences appeared months or years beforehand in large numbers of chronic-infection sequences. For example, the most common mutations that we find in our EPCI dataset through mid-2022 include S:ΔY144, S:R346T, S:N460K, S:R493Q, ORF1a:L3829F, ORF1a:T4175I, and many others that, at some later time period, became nearly universal in circulating viral genomes. The extremely high rate of ORF9b mutations during chronic infections—higher than in any other viral protein relative to its size (Figure 3B)—along with the striking convergence exhibited in these mutations (Supplementary table S3), suggest ORF9b is subject to intense selection pressures in the context of long-term infections.

**Figure 3:**
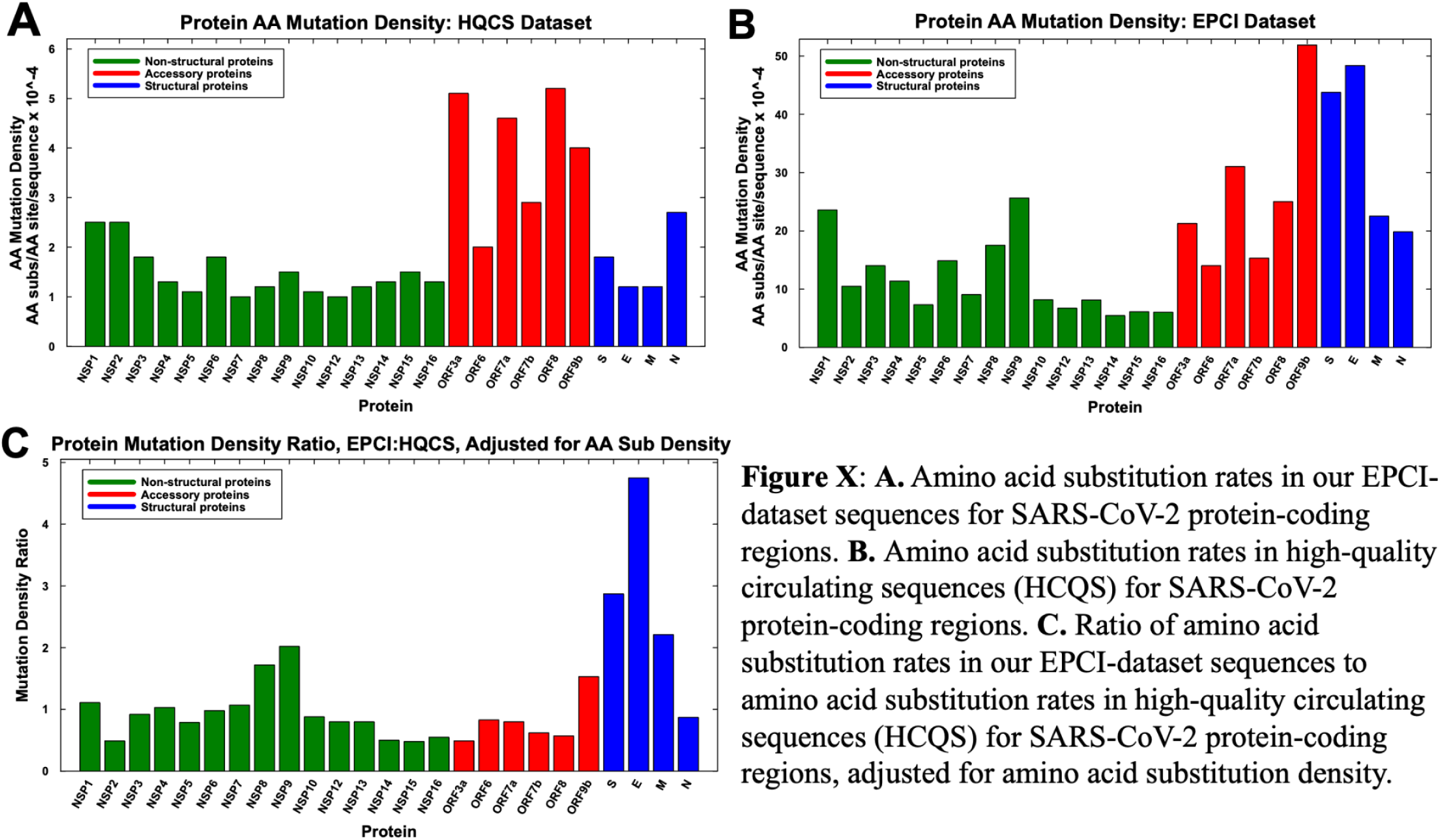
**A.** Amino acid substitution rates in our EPCI-dataset sequences for SARS-CoV-2 protein-coding regions. **B.** Amino acid substitution rates in high-quality circulating sequences (HCQS) for SARS-CoV-2 protein-coding regions. **C.** Ratio of amino acid substitution rates in our EPCI-dataset sequences to amino acid substitution rates in high-quality circulating sequences (HCQS) for SARS-CoV-2 protein-coding regions.

We offer three non-exclusive explanations for why the frequencies of mutations in chronic infection-derived sequences (i.e. sequences enriched for in our EPCI dataset) might differ from those seen in circulating sequences (i.e. those in our HQCS dataset). First, intra-host evolution may select for traits that facilitate long-term infection in various ways but which impair interhost transmission. Deletions in the spike NTD, for example, have been shown to increase infectivity^78–79^ (refs), which could favor long-term survival within a host. But these deletions also increase instability and premature S1 shedding, which may decrease the time a virus remains infectious within aerosols, thereby inhibiting transmission^78^.

Second, most long-term infections involve immunocompromised hosts. Immune-system-imposed evolutionary constraints operating in immunocompetent hosts are likely to be relaxed in the immunocompromised, permitting mutations that are deleterious in immunocompetent humans to occur. Because there are myriad ways to be immunocompromised, each having distinct effects on the complex, multifaceted human immune system, a multitude of intra-host selection regimes must exist in our world of over 8 billion people, giving rise to a variety of mutations impeding infection of immunocompetent hosts.

Finally, the absence of a transmission bottleneck during chronic infection leads to greatly accelerated adaptive evolution^74^ (refs). The stochastic element introduced by the very tight SARS-CoV-2 transmission bottleneck, together with the brief time periods between infection and transmission, likely prevent selection of many modestly advantageous mutations that, during long-term infection, can gradually increase in prevalence over months or years of uninterrupted intra-host evolution^80–81^ (refs).

### Chronic-infection derived sequences display unusually high frequencies of amino acid substitutions in ORF9b

In circulating sequences (i.e. our HQCS dataset) ORF9b does not display an unusually elevated density of size-adjusted amino acid substitutions relative to other accessory protein genes (note red bars in Figure 3A). In EPCI sequences, however, ORF9b clearly displays the highest density of substitutions of any gene (note red bars in Figure 3B), and is the accessory protein with the highest disparity of substitution frequencies relative to HQCS sequences (note red bars in Figure 3C; Supplementary Table S2).

Looking more closely at the distribution of amino acid substitutions in ORF9b (Figure 4 and Supplementary Figure X) it is evident that the disparities in EPCI and HQCS substitution frequencies in ORF9b map primarily to the 17 N-terminal and 12 C-terminal amino acid sites of the protein;(and secondarily to the region around residue 64.

**Figure 4:**
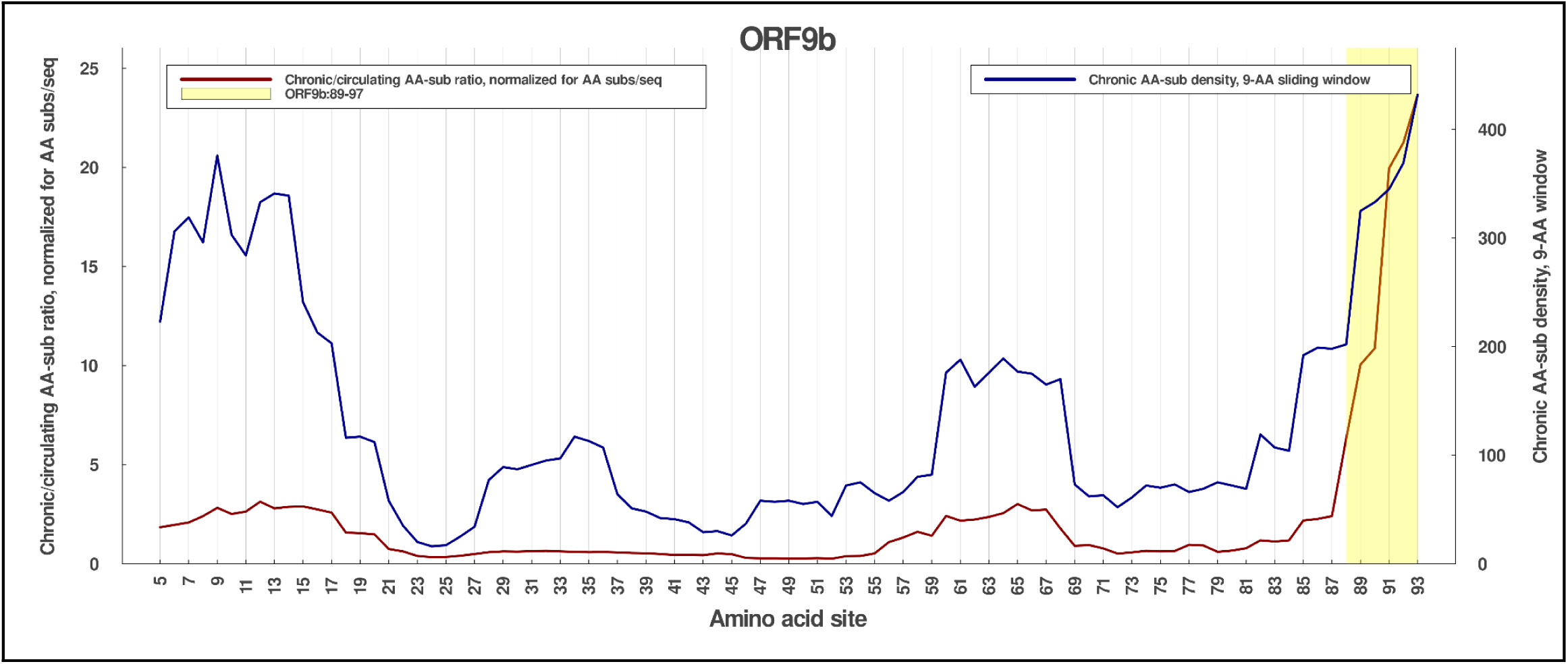
The total number of ORF9b substitutions in each nine-amino acid sliding window in the EPCI dataset (blue) and the ratio of ORF9b substitutions per sequence in our EPCI dataset to ORF9b substitutions in high-quality circulating sequences, adjusted for the number of amino acid substitutions per EPCI sequence relative to high-quality circulating sequences (red).

**Figure 5:**
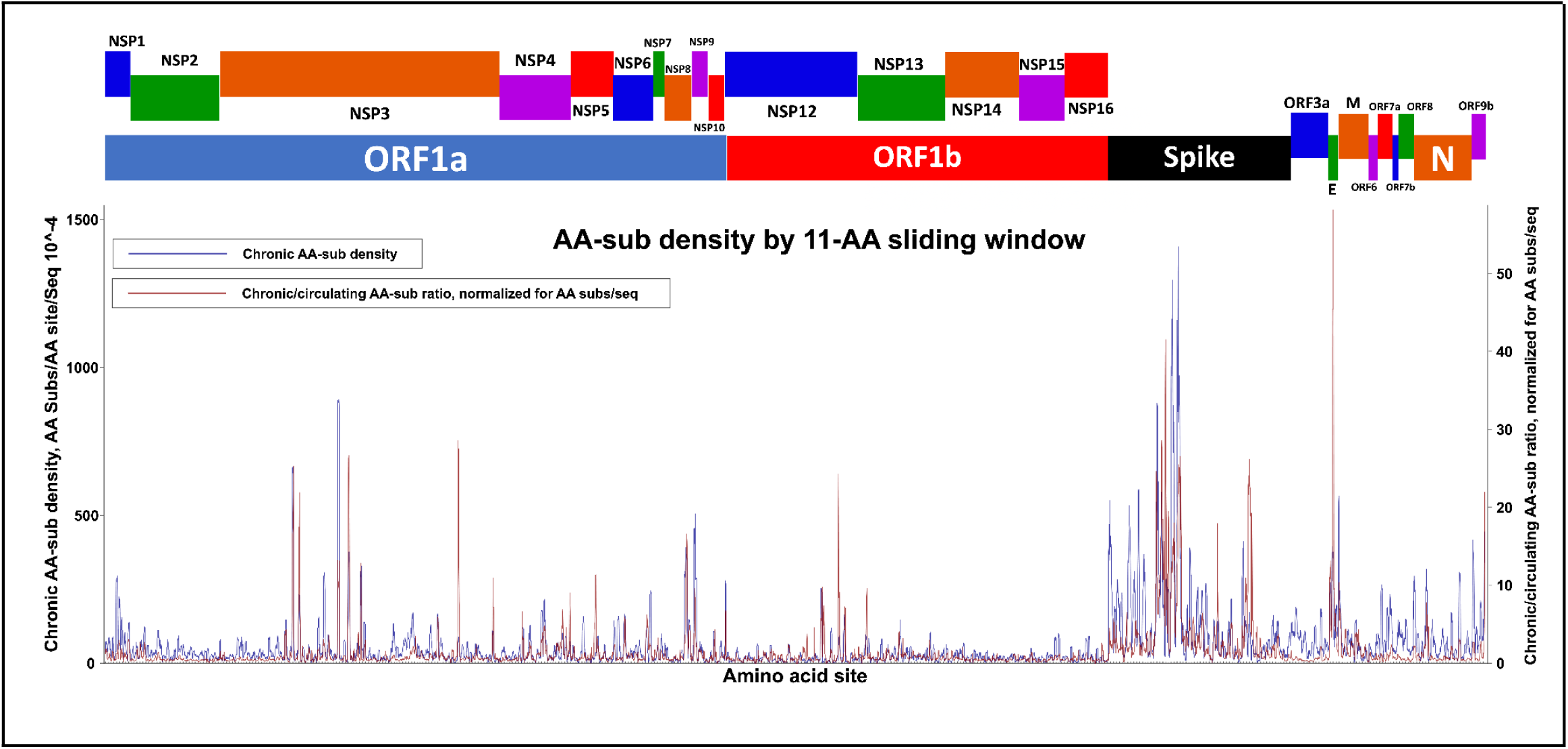
The total number of amino acid substitutions in each 11-amino acid sliding window for each protein in the SARS-CoV-2 genome in our EPCI dataset (blue) and the ratio of amino acid substitutions per sequence in the EPCI dataset to amino acid substitutions in high-quality circulating sequences, adjusted for the number of amino acid substitutions per EPCI sequence relative to high-quality circulating sequences (red).

### During chronic infections ORF9b N-terminal substitutions tend to converge on states seen in other sarbecoviruses

Some of the ORF9b N-terminal domain (NTD) substitution that are among the most enriched in the EPCI dataset relative to the HQCS dataset, such as ORF9b:I5T (4.18-fold enrichment), P10S (7.39-fold), F10S (93.03), A11E (161.02-fold), and R13H (23.95-fold). Two of these have also been found in many of the most successful circulating variants (e.g. P10S is also found in all major Omicron lineages and I5T in XBB.1.9 lineages), suggesting that they confer a selective advantage both within persistently infected hosts and in circulation (Supplementary Table S3).

XBB lineages emerged in December 2022 possessing the key S:S486P mutation, which dramatically increased spike ACE2 affinity and conferred a powerful growth advantage. S:S486P first appeared in XBB.1.5, but soon after XBB.1.16, XBB.1.9.1, XB.1.9.2 emerged and decisively outcompeted XBB.1.5 despite having identical spikes. The only amino acid mutation shared by these three fitter lineages was ORF9b:I5T, indicating it alone likely conferred a distinct competitive advantage, at least in the context of an XBB genetic background. ORF9b:I5T is a change to the AA residue found in other sarbecoviruses including SARS-CoV and close SARS-CoV-2 relatives sampled from bats (such as RaTG13). Other ORF9b substitutions that are notably enriched (from 3.24- to 43.61-fold) among our EPCI sequences and result in “reversion” to amino acid states found in other sarbecoviruses include M8V, R13H, P51Q, N62D, K67R, L71S, and V96A: a pattern suggestive of either adaptation to the gastrointestinal tract—believed to be the major target of infection in bat coronaviruses^86^—or adaptation for long-term within-host persistence, as is also thought to occur in bats^91^.

### High substitution frequencies in the C-terminal domain of ORF9b during chronic infections are potentially driven by immune pressures

Another distinctive ORF9b mutational pattern that we find is clusters of two or three mutations occurring significantly more commonly (p = <0.00001, Fisher exact test) in the last nine c-terminal domain (CTD) residues of ORF9b of individual EPCI dataset sequences than in individual HQCSs. Three of the most common private mutations seen in our EPCI sequences (frequency > 1.8%) are D89E (70.85-fold enrichment), V93L (133.37-fold enrichment), and K97E (516.34-fold enrichment), all of which very rarely occur in HQCS lineages (frequency <0.08%). The only individual HQCS lineage that provides a minor exception to this trend is BA.2.3.20 which carries an ORF9b:D89E mutation and accounts for 85% of HQCSs with either D89E, V93L, or K97E. This lineage likely originated in the Philippines where it was the dominant variant for several months in 2022. It is, however, noteworthy that the BA.2.3.20 clade is separated from the rest of the SARS-CoV-2 tree by an extremely long-branch, likely indicating a chronic-infection origin.

The high density of EPCI enriched mutations between ORF9b sites 89-97 are all the more notable for their occurrence in codons encoding a particularly essential region of N: the RNA-binding finger located at N:90-106, a highly conserved, essential region for N function^62^. The nucleotide substitution resulting in ORF9b:D89E is synonymous in N while that resulting in ORF9b:V93L/M results in either N:G96V, N:G96A, or N:G96D. The nucleotide substitution resulting in ORF9b:K97E corresponds also results in the conservative N:K100R mutation. ORF9b:K97E is among the most distinctive chronic-infection mutations, arising independently 527 times more frequently in EPCI sequences than in HQCS sequences (after normalizing for the greater number of private mutations in EPCI sequences).

ORF9b:89-97 is exposed and disordered in the ORF9b-mon structure. This raises the possibility that residues 89-97 could be either a target of antibodies or a T-cell epitope. As non-spike-targeting antibodies are not expected to be neutralizing, they are unlikely to have a strong effect on transmission in circulating lineages. They may, however, be important for viral clearance, and therefore mutations that evade such immune responses could be subject to positive selection during chronic infections. Furthermore, the ORF9b CTD is likely ubiquitinated, which leads to proteasomal degradation^82^, hinting that ORF9b CTD mutations may interrupt this degradation, maintaining more robust, longer-term innate immune antagonism.

While nearly all studies of SARS-CoV-2 T cell epitopes neglect ORF9b, one study of non-canonical SARS-CoV-2 ORFs found that the ORF9b:89-96 peptide (ELPDEFVVVTV) provoked the strongest CD8+ T cell response among all peptides tested in seven convalescent humans expressing HLA-A*02:01^88^. Notably, the SLEDKAFQL peptide, spanning ORF9b:63-71, also ranked among the top five most reactive epitopes tested in this study^88^. Mutations in these two regions often appear together in EPCI sequences, suggesting selection for T cell evasion may be one cause for these ORF9b mutations.

The other major exposed, disordered region in both the ORF9b-mon and ORF9b-dim structures is the N-terminal domain, where ORF9b:I5T, the most common ORF9b mutation in our EPCI dataset outside the CTD (Supplementary table S3), is found. The 4.11 times higher frequency of independent development of this mutation in the EPCI dataset relative to HQCS dataset is consistent with the possibility that I5T is an immune-evasion mutation.

### Mutations that potentially impact the affinity with which ORF9b-mon binds to TOM70 are overrepresented in chronic infection derived sequences

Purified ORF9b protein is dominated by the ORF9b-dim form, which is unable to bind to TOM70. The formation of ORF9b-mon only occurs in the presence of cytoplasmic factors, as when coexpressed with TOM70 in cells or when added to a TOM70-containing rabbit reticulocyte lysate. Under these conditions, ORF9b-mon forms complexes similar in size to those formed by chaperone proteins and mitochondrial pre-proteins, suggesting that, after translation, ORF9b-mon might be held in its monomer form and delivered to the mitochondrial membrane by cellular chaperones^10^.

ORF9b-mon, consists of a central region featuring a long central alpha helix surrounded by disordered regions. The helical ORF9b-mon 43-76 region binds tightly to the TOM70 CTD pocket primarily through hydrophobic interactions, driving a conformational change that causes a 29-fold reduction in HSP90 affinity for the TOM70 NTD^6^. The delivery of TBK1 and IRF3 is therefore prevented, interrupting the RIG-I-initiated signaling cascade and efficiently antagonizing the production of type-I IFN.

The TOM70 CTD pocket binds to the mitochondrial targeting signal (MTS) of chaperone-associated mitochondrial pre-proteins, and the central ORF9b-mon residues that bind this region have been shown to likely contain an MTS^7^. The TOM70-ORF9b-mon association is dominated by hydrophobic interactions. Crucial to the TOM70-CTD-ORF9b-mon interaction is hydrogen bonding between TOM70:E477 and the highly conserved ORF9b:S53 residue^6^.

ORF9b:S53 is one of two sites in the ORF9b-mon 43-76 region (the other being ORF9b:S50) that may be phosphorylated at some point during an infection cycle. Phosphorylation of the highly conserved ORF9b:S53 powerfully reduces binding to TOM70^10, 38^, suggesting phosphorylation may regulate a shift in ORF9b form and function between early and late infection, likely from ORF9b-mon to ORF9b-dim.

The Alpha variant, which featured the strongest innate immune evasion of any early VOC lineage, showed less phosphorylation of ORF9b early in infection and higher rates of phosphorylation late in infection, an adaptation that may have improved ORF9b’s ability to both suppress IFN production and assist in viral assembly^10, 38^.

The huge disparity in the degree of TOM70-ORF9b-mon affinity reduction observed between the central, alpha-helix region of ORF9b-mon (residues 43-78) and full-length ORF9b-mon is one of several indications that ORF9b-mon binds to HSP90 in two locations; in addition to the well described binding to the TOM70 CTD by ORF9b:43-78, a second binding site in the TOM70 NTD likely exists. This site corresponds to the positively charged NTD-pocket to which HSP90 and other chaperone proteins bind primarily through hydrophilic interactions^38^.

A key TOM70 residue for the binding of chaperone proteins to the TOM70 NTD site is R192. Specifically, in an R192A mutant, ORF9b binding to TOM70 was reduced by 50%, suggesting that interaction with the TOM70 NTD is a vital part of the ORF9b-TOM70 association^38^.

The ORF9b-mon CTD is disordered and is not represented in any experimentally determined ORF9b structures, but, as Gao et al note^6^, it is located in the same general region where the HSP90 EEVD motif binds to TOM70. The ORF9b-mon CTD has an EELPDE motif from 85-90 that is highly conserved both within SARS-CoV-2 and across related sarbecoviruses. By far the most common mutations in this region in both circulating and chronic SARS-CoV-2 sequences are E86D and D89E, both conservative mutations that maintain negative charge. Might the negatively charged ORF9b:85-90 region bind, like HSP90’s EEVD motif, to the TOM70 NTD, increasing ORF9b’s affinity for TOM70 and inhibiting HSP90 binding? Some of the mutations seen in chronic-infection sequences in this region also hint that this might be the case.

Specifically, mutations that increase the negative charge of ORF9b residues 85 to 90 such as ORF9b:K97E (519.6-fold increase in EPCI dataset relative to HQCS dataset; Supplementary Table S3) and ORF9b:V96E (a 34.98-fold increase in EPCI; SUpplementary Table S3) may be favoured because they increase the affinity of ORF9b-mon for TOM70.

Two other ORF9b:85-90 mutations that are overrepresented in our EPCI dataset are ORF9b:D89E (63.81-fold increase), and V93L (118.2-fold increase). Although both mutations are conservative with respect to maintaining the negative charge of this region, it is still plausible that they might impact the affinity with which ORF9b-mon binds TOM70.

### The ORF1a:K1795Q mutation that is most characteristic of sequences derived from chronic infections potentially modulates the ubiquitination/deubiquitination of ORF9b

In addition to phosphorylation, ubiquitination also regulates ORF9b activity. Ubiquitin is a small, cellular protein that regulates a multitude of cellular and viral processes, including the antiviral response^83^. Ubiquitin chains are formed by covalent linkage of the C-terminal glycine of one ubiquitin molecule to one of the seven internal lysine (K) residues of another ubiquitin molecule, a reaction catalyzed by a variety of cellular E3 ubiquitin ligases. Ubiquitin chains linked through the K48 residue typically mark target proteins for degradation by the proteasome, which recycles the resulting protein components for reuse. Of the eight SARS-CoV-2 proteins that have been experimentally assessed in this regard, only ORF9b was found to undergo ubiquitination-mediated proteasomal degradation^82^.

Three cellular E3 ubiquitin ligases were found to be responsible for K48-linked ubiquitination of ORF9b^84^, while the cellular deubiquitinase USP29 was found to prevent proteasomal degradation of ORF9b, leading to an ORF9b-mediated increase in virulence in cells infected with either vesicular stomatitis virus (VSV) or transcription and replication-competent SARS-CoV-2 virus-like particles (trVLP). Furthermore, USP29 mRNA levels are increased in the peripheral blood mononuclear cells of SARS-CoV-2-infected patients compared to healthy controls, suggesting that prevention or reversal of ORF9b ubiquitination could possibly play a role in COVID-19 pathogenesis^82^.

Intriguingly, USP29 was found to bind to ORF9b primarily through the ORF9b C-terminus (residues 68-97), which we have found to be one of the most intensely mutated regions of the entire SARS-CoV-2 genome in our EPCI sequences (Supplementary Figure X). Given the frequency with which mutations thought to increase ORF9b expression occur in long-term SARS-CoV-2 infections, it would be interesting to test whether the numerous highly convergent ORF9b CTD mutations in sequences derived from likely chronic infections affect ORF9b-USP29 binding strength.

Speculatively, the vulnerability of ORF9b to ubiquitination and degradation may partially explain why the Gamma VOC was one of only two VOI/VOC lineages to lack a mutation reducing the Kozak-sequence favorability of the N start codon. Gamma was the only major SARS-CoV-2 lineage to possess the ORF1a:K1795Q mutation, which has been shown to dramatically increase the ability of the NSP3 papain-like protease (PLpro) to remove K48-linked ubiquitin from its targets^85^. It seems plausible, therefore, that the PLpro of Gamma acts to prevent ORF9b degradation in a similar way to the cellular ubiquitinase, USP29. Conversely, *all* sarbecoviruses possess a glutamine residue at the PLpro site corresponding to ORF1a:1795 as well as favorable N Kozak sequences.

There are indications that in bats, coronaviruses preferentially infect the gastrointestinal tract^86^. The common presence of ORF1a:K1795Q in cryptic wastewater lineages—which evidence strongly suggests originate from very long-term infections in individual humans—and in bat sarbecoviruses, along with its absence in virtually all SARS-CoV-2 lineages throughout the pandemic (Gamma being the only notable exception) may suggest that ORF1a:K1795Q is a favorable mutation in the GI tract but is deleterious for respiratory transmission^87^. If this is true, the now-universal N Kozak-sequence mutations shown to increase ORF9b protein expression may be a way of compensating for increased proteasomal degradation of ORF9b in transmitting SARS-CoV-2 variants that lack the ORF1a:K1795Q mutation found in all other known sarbecoviruses.

## Conclusion

The dynamic and convergent evolution of ORF9b and genome regions determining its expression levels across multiple diverse SARS-CoV-2 lineages throughout the COVID-19 pandemic, indicates the vital, albeit underappreciated, role this protein has played as the virus has adapted to infecting humans, particularly after vaccination and multiple waves of infection have granted much of the human population some degree of immunity. The high substitution rate in ORF9b sequences sampled from individuals with likely chronic infections suggests that ORF9b-mon and/or ORF-9b-dim may play important roles in viral persistence and immune evasion. Whereas ORF-9b-mon likely mediates suppression of host type-I IFN responses, the replication defect seen in IFN-suppressed primary human cells hints that it likely has at least one other presently unknown role in the viral life cycle. Further, evidence drawn from other betacoronaviruses suggests that ORF9b-dim could be a structural component of virions and may contribute to viral assembly. Although the sub-cellular localization and function of ORF9b-mon has been studied in some detail, it remains unknown where ORF9b-dim is localized in the cell and what role it plays during infection. Given that intense selection on ORF9b expression levels was likely a central feature of SARS-CoV-2 evolution during the first two years of the pandemic, and that adaptive evolution of ORF9b is likely ongoing, especially in the context of chronic infections, a more complete determination of ORF9b functions should be a priority of future SARS-CoV-2 molecular virology research.

## Additional Information

The custom Julia code used for this paper can be found at https://github.com/ryhisner/ORF9b_relevant_code. GISAID accession numbers for the sequences in the EPCI dataset can be found at https://github.com/ryhisner/EPCI_dataset.

## Acknowledgements

We gratefully acknowledge all data contributors, i.e., the Authors and their Originating laboratories responsible for obtaining the specimens, and their Submitting laboratories for generating the genetic sequence and metadata and sharing via the GISAID Initiative, on which this research is based.

## Notes

### Competing Interest Statement

The authors have declared no competing interest.

## References

1. Wong LR, Perlman S. Immune dysregulation and immunopathology induced by SARS-CoV-2 and related coronaviruses - are we our own worst enemy? [published correction appears in Nat Rev Immunol. 2022 Mar;22(3):200. doi: 10.1038/s41577-021-00673-1.]. Nat Rev Immunol. 2022;22(1):47–56. doi:10.1038/s41577-021-00656-2

2. Lucas C, Wong P, Klein J, et al. Longitudinal analyses reveal immunological misfiring in severe COVID-19. Nature. 2020;584(7821):463–469. doi:10.1038/s41586-020-2588-y

3. Hadjadj J, Yatim N, Barnabei L, et al. Impaired type I interferon activity and inflammatory responses in severe COVID-19 patients. Science. 2020;369(6504):718–724. doi:10.1126/science.abc6027

4. Meier C, Aricescu AR, Assenberg R, et al. The crystal structure of ORF-9b, a lipid binding protein from the SARS coronavirus. Structure. 2006;14(7):1157–1165. doi:10.1016/j.str.2006.05.012

5. Jin X, Sun X, Chai Y, et al. Structural characterization of SARS-CoV-2 dimeric ORF9b reveals potential fold-switching trigger mechanism. Sci China Life Sci. 2023;66(1):152–164. doi:10.1007/s11427-022-2168-8

6. Gao X, Zhu K, Qin B, Olieric V, Wang M, Cui S. Crystal structure of SARS-CoV-2 Orf9b in complex with human TOM70 suggests unusual virus-host interactions. Nat Commun. 2021;12(1):2843. Published 2021 May 14. Doi:10.1038/s41467-021-23118-8

7. Gordon DE, Hiatt J, Bouhaddou M, et al. Comparative host-coronavirus protein interaction networks reveal pan-viral disease mechanisms. Science. 2020;370(6521):eabe9403. doi:10.1126/science.abe9403

8. Lowery SA, Schuster N, Wong L-YR, et al. Mouse hepatitis virus JHMV I protein is required for maximal virulence. J Virol. 2024;98(9):e0068024. doi:10.1128/jvi.00680-24

9. Xu K, Zheng BJ, Zeng R, et al. Severe acute respiratory syndrome coronavirus accessory protein 9b is a virion-associated protein. Virology. 2009;388(2):279–285. doi:10.1016/j.virol.2009.03.032

10. Brandherm L, Kobaš AM, Klöhn M, et al. Phosphorylation of SARS-CoV-2 Orf9b Regulates Its Targeting to Two Binding Sites in TOM70 and Recruitment of Hsp90. Int J Mol Sci. 2021;22(17):9233. Published 2021 Aug 26. doi:10.3390/ijms22179233

11. Jiang HW, Zhang HN, Meng QF, et al. SARS-CoV-2 Orf9b suppresses type I interferon responses by targeting TOM70. Cell Mol Immunol. 2020;17(9):998–1000. doi:10.1038/s41423-020-0514-8

12. Fischer F, Peng D, Hingley ST, Weiss SR, Masters PS. The internal open reading frame within the nucleocapsid gene of mouse hepatitis virus encodes a structural protein that is not essential for viral replication. J Virol. 1997;71(2):996–1003. doi:10.1128/JVI.71.2.996-1003.1997

13. Li Y, Xu Z, Lei Q, et al. Antibody landscape against SARS-CoV-2 reveals significant differences between non-structural/accessory and structural proteins. Cell Rep. 2021;36(2):109391. doi:10.1016/j.celrep.2021.109391

14. Jiang HW, Li Y, Zhang HN, et al. SARS-CoV-2 proteome microarray for global profiling of COVID-19 specific IgG and IgM responses. Nat Commun. 2020;11(1):3581. Published 2020 Jul 14. doi:10.1038/s41467-020-17488-8

15. Qiu M, Shi Y, Guo Z, et al. Antibody responses to individual proteins of SARS coronavirus and their neutralization activities. Microbes Infect. 2005;7(5-6):882–889. doi:10.1016/j.micinf.2005.02.006

16. Senanayake SD, Hofmann MA, Maki JL, Brian DA. The nucleocapsid protein gene of bovine coronavirus is bicistronic. J Virol. 1992;66(9):5277–5283. doi:10.1128/JVI.66.9.5277-5283.1992

17. Khare, S., et al (2021) GISAID’s Role in Pandemic Response. China CDC Weekly, 3(49): 1049–1051. doi: 10.46234/ccdcw2021.255

18. Aksamentov, I., Roemer, C., Hodcroft, E. B., & Neher, R. A., (2021). Nextclade: clade assignment, mutation calling and quality control for viral genomes. Journal of Open Source Software, 6(67), 3773, 10.21105/joss.03773 https://clades.nextstrain.org/

19. Chen, C., Nadeau, S., Yared, M., Voinov, P., Ning, X., Roemer, C. & Stadler, T. “CoV-Spectrum: Analysis of globally shared SARS-CoV-2 data to Identify and Characterize New Variants” Bioinformatics (2021); doi: 10.1093/bioinformatics/btab856. https://cov-spectrum.org/

20. Turakhia Y, Thornlow B, Hinrichs AS, et al. Ultrafast Sample placement on Existing tRees (UShER) enables real-time phylogenetics for the SARS-CoV-2 pandemic. Nat Genet. 2021;53(6):809–816. doi:10.1038/s41588-021-00862-7 https://genome.ucsc.edu/cgi-bin/hgPhyloPlace

21. Roch S, Nute M, Warnow T. Long-Branch Attraction in Species Tree Estimation: Inconsistency of Partitioned Likelihood and Topology-Based Summary Methods. Syst Biol. 2019;68(2):281–297. doi:10.1093/sysbio/syy061

22. Meng EC, Goddard TD, Pettersen EF, et al. UCSF ChimeraX: Tools for structure building and analysis. Protein Sci. 2023;32(11):e4792. doi:10.1002/pro.4792

23. Li Y, Jin Y, Kuang L, et al. The N-Terminal Region of Middle East Respiratory Syndrome Coronavirus Accessory Protein 8b Is Essential for Enhanced Virulence of an Attenuated Murine Coronavirus. J Virol. 2022;96(3):e0184221. doi:10.1128/JVI.01842-21

24. Simon-Loriere E, Holmes EC, Pagán I. The effect of gene overlapping on the rate of RNA virus evolution. Mol Biol Evol. 2013;30(8):1916–1928. doi:10.1093/molbev/mst094

25. Sawicki SG, Sawicki DL, Siddell SG. A contemporary view of coronavirus transcription. J Virol. 2007;81(1):20–29. doi:10.1128/JVI.01358-06

26. Sola I, Almazán F, Zúñiga S, Enjuanes L. Continuous and Discontinuous RNA Synthesis in Coronaviruses. Annu Rev Virol. 2015;2(1):265–288. doi:10.1146/annurev-virology-100114-055218

27. Lan TCT, Allan MF, Malsick LE, et al. Secondary structural ensembles of the SARS-CoV-2 RNA genome in infected cells. Nat Commun. 2022;13(1):1128. Published 2022 Mar 2. doi:10.1038/s41467-022-28603-2

28. Finkel Y, Mizrahi O, Nachshon A, et al. The coding capacity of SARS-CoV-2. Nature. 2021;589(7840):125–130. doi:10.1038/s41586-020-2739-1

29. Cohen P, DeGrace EJ, Danziger O, Patel RS, Barrall EA, Bobrowski T, Kehrer T, Cupic A, Miorin L, García-Sastre A, Rosenberg BR.2023.Unambiguous detection of SARS-CoV-2 subgenomic mRNAs with single-cell RNA sequencing. Microbiol Spectr11:e00776–23.10.1128/spectrum.00776-23

30. Wang D, Jiang A, Feng J, et al. The SARS-CoV-2 subgenome landscape and its novel regulatory features. Mol Cell. 2021;81(10):2135–2147.e5. doi:10.1016/j.molcel.2021.02.036

31. Parker MD, Stewart H, Shehata OM, et al. Altered subgenomic RNA abundance provides unique insight into SARS-CoV-2 B.1.1.7/Alpha variant infections. Commun Biol. 2022;5(1):666. Published 2022 Jul 5. doi:10.1038/s42003-022-03565-9

32. Wong L-YR, Odle A, Luhmann E, et al. Contrasting roles of MERS-CoV and SARS-CoV-2 internal proteins in pathogenesis in mice. mBio. 2023;14(6):e0247623. doi:10.1128/mbio.02476-23

33. Lee JS, Dittmar M, Miller J, et al. Pressure to evade cell-autonomous innate sensing reveals interplay between mitophagy, IFN signaling, and SARS-CoV-2 evolution. Cell Rep. 2025;44(1):115115. doi:10.1016/j.celrep.2024.115115

34. Kozak M. Pushing the limits of the scanning mechanism for initiation of translation. Gene. 2002;299(1-2):1–34. doi:10.1016/s0378-1119(02)01056-9

35. Kozak M. Point mutations define a sequence flanking the AUG initiator codon that modulates translation by eukaryotic ribosomes. Cell. 1986;44(2):283–292. doi:10.1016/0092-8674(86)90762-2

36. Wang J, Peng Y, Zhao L, Cao M, Hung T, Deng T. Influenza A virus utilizes a suboptimal Kozak sequence to fine-tune virus replication and host response. J Gen Virol. 2015;96(Pt 4):756–766. doi:10.1099/vir.0.000030

37. Shukla N, Kamath ND, Snell JC, Bruchez AM, Matreyek KA. Kozak sequence libraries for characterizing transgenes across expression levels. Published online April 30, 2025. 10.1101/2025.04.28.651141

38. Thorne, L.G., Bouhaddou, M., Reuschl, AK. et al. Evolution of enhanced innate immune evasion by SARS-CoV-2. Nature 602, 487–495 (2022). 10.1038/s41586-021-04352-y

39. Reuschl AK, Thorne LG, Whelan MVX, et al. Evolution of enhanced innate immune suppression by SARS-CoV-2 Omicron subvariants. Nat Microbiol. 2024;9(2):451–463. doi:10.1038/s41564-023-01588-4

40. Xie J, Zhuang Z, Gou S, et al. Precise genome editing of the Kozak sequence enables bidirectional and quantitative modulation of protein translation to anticipated levels without affecting transcription. Nucleic Acids Res. 2023;51(18):10075–10093. doi:10.1093/nar/gkad687

41. Wu H, Xing N, Meng K, et al. Nucleocapsid mutations R203K/G204R increase the infectivity, fitness, and virulence of SARS-CoV-2. Cell Host Microbe. 2021;29(12):1788–1801.e6. doi:10.1016/j.chom.2021.11.005

42. Mears HV, Young GR, Sanderson T, et al. Emergence of SARS-CoV-2 subgenomic RNAs that enhance viral fitness and immune evasion. PLoS Biol. 2025;23(1):e3002982. Published 2025 Jan 21. doi:10.1371/journal.pbio.3002982

43. Ricciardi S, Guarino AM, Giaquinto L, et al. The role of NSP6 in the biogenesis of the SARS-CoV-2 replication organelle. Nature. 2022;606(7915):761–768. doi:10.1038/s41586-022-04835-6

44. Senanayake SD, Brian DA. Bovine coronavirus I protein synthesis follows ribosomal scanning on the bicistronic N mRNA. Virus Res. 1997;48(1):101–105. doi:10.1016/s0168-1702(96)01423-2

45. Onomoto K, Onoguchi K, Yoneyama M. Regulation of RIG-I-like receptor-mediated signaling: interaction between host and viral factors. Cell Mol Immunol. 2021;18(3):539–555. doi:10.1038/s41423-020-00602-7

46. Hou F, Sun L, Zheng H, Skaug B, Jiang QX, Chen ZJ. MAVS forms functional prion-like aggregates to activate and propagate antiviral innate immune response [published correction appears in Cell. 2011 Sep 2;146(5):841]. Cell. 2011;146(3):448–461. doi:10.1016/j.cell.2011.06.041

47. Mishra S, Looger LL, Porter LL. A sequence-based method for predicting extant fold switchers that undergo α-helix ↔ β-strand transitions. Biopolymers. 2021;112(10):e23471. doi:10.1002/bip.23471

48. San Felipe CJ, Batra J, Muralidharan M, et al. Coupled equilibria of dimerization and lipid binding modulate SARS Cov 2 Orf9b interactions and interferon response. Elife. 2025;14:RP106484. Published 2025 Sep 17. doi:10.7554/eLife.106484

49. Zhai, Q., Fisher, R., Chung, HY. et al. Structural and functional studies of ALIX interactions with YPXnL late domains of HIV-1 and EIAV. Nat Struct Mol Biol 15, 43–49 (2008). 10.1038/nsmb1319

50. Zhang Y, Huang L, Ren C, Wang W, Wang X, Gao G. β-Coronaviruses exploit ESCRT for virion assembly and egress. mBio. Published online May 23, 2025. doi:10.1128/mbio.00979-25

51. Barr JN, Wertz GW. Polymerase slippage at vesicular stomatitis virus gene junctions to generate poly(A) is regulated by the upstream 3’-AUAC-5’ tetranucleotide: implications for the mechanism of transcription termination. J Virol. 2001;75(15):6901–6913. doi:10.1128/JVI.75.15.6901-6913.2001

52. Morita E, Sandrin V, McCullough J, Katsuyama A, Baci Hamilton I, Sundquist WI. ESCRT-III protein requirements for HIV-1 budding. Cell Host Microbe. 2011;9(3):235–242. doi:10.1016/j.chom.2011.02.004

53. Strack B, Calistri A, Craig S, Popova E, Göttlinger HG. AIP1/ALIX is a binding partner for HIV-1 p6 and EIAV p9 functioning in virus budding. Cell. 2003;114(6):689–699. doi:10.1016/s0092-8674(03)00653-6

54. Barouch-Bentov R, Neveu G, Xiao F, et al. Hepatitis C Virus Proteins Interact with the Endosomal Sorting Complex Required for Transport (ESCRT) Machinery via Ubiquitination To Facilitate Viral Envelopment [published correction appears in mBio. 2018 Jan 9;9(1):e02234-17. doi: 10.1128/mBio.02234-17.]. mBio. 2016;7(6):e01456-16. Published 2016 Nov 1. doi:10.1128/mBio.01456-16

55. Shirasaki T, Feng H, Duyvesteyn HME, et al. Nonlytic cellular release of hepatitis A virus requires dual capsid recruitment of the ESCRT-associated Bro1 domain proteins HD-PTP and ALIX. PLoS Pathog. 2022;18(8):e1010543. Published 2022 Aug 15. doi:10.1371/journal.ppat.1010543

56. Aloise C, Schipper JG, van Vliet A, et al. SARS-CoV-2 nucleocapsid protein inhibits the PKR-mediated integrated stress response through RNA-binding domain N2b. PLoS Pathog. 2023;19(8):e1011582. Published 2023 Aug 22. doi:10.1371/journal.ppat.1011582

57. Nabeel-Shah S, Lee H, Ahmed N, et al. SARS-CoV-2 nucleocapsid protein binds host mRNAs and attenuates stress granules to impair host stress response. iScience. 2022;25(1):103562. doi:10.1016/j.isci.2021.103562

58. Lu S, Ye Q, Singh D, et al. The SARS-CoV-2 nucleocapsid phosphoprotein forms mutually exclusive condensates with RNA and the membrane-associated M protein. Nat Commun. 2021;12(1):502. Published 2021 Jan 21. doi:10.1038/s41467-020-20768-y

59. Bessa LM, Guseva S, Camacho-Zarco AR, et al. The intrinsically disordered SARS-CoV-2 nucleoprotein in dynamic complex with its viral partner nsp3a. Sci Adv. 2022;8(3):eabm4034. doi:10.1126/sciadv.abm4034

60. Scherer KM, Mascheroni L, Carnell GW, et al. SARS-CoV-2 nucleocapsid protein adheres to replication organelles before viral assembly at the Golgi/ERGIC and lysosome-mediated egress. Sci Adv. 2022;8(1):eabl4895. doi:10.1126/sciadv.abl4895

61. Yang Z, Johnson BA, Meliopoulos VA, et al. Interaction between host G3BP and viral nucleocapsid protein regulates SARS-CoV-2 replication and pathogenicity. Cell Rep. 2024;43(3):113965. doi:10.1016/j.celrep.2024.113965

62. Botova M, Camacho-Zarco AR, Tognetti J, et al. A specific phosphorylation-dependent conformational switch in SARS-CoV-2 nucleocapsid protein inhibits RNA binding. Sci Adv. 2024;10(31):eaax2323. doi:10.1126/sciadv.aax2323

63. Syed AM, Ciling A, Chen IP, et al. SARS-CoV-2 evolution balances conflicting roles of N protein phosphorylation. PLoS Pathog. 2024;20(11):e1012741. Published 2024 Nov 21. doi:10.1371/journal.ppat.1012741

64. Carlson CR, Asfaha JB, Ghent CM, et al. Phosphoregulation of Phase Separation by the SARS-CoV-2 N Protein Suggests a Biophysical Basis for its Dual Functions. Mol Cell. 2020;80(6):1092–1103.e4. doi:10.1016/j.molcel.2020.11.025

65. Carlson CR, Adly AN, Bi M, et al. Reconstitution of the SARS-CoV-2 ribonucleosome provides insights into genomic RNA packaging and regulation by phosphorylation. J Biol Chem. 2022;298(11):102560. doi:10.1016/j.jbc.2022.102560

66. Chaguza C, Hahn AM, Petrone ME, et al. Accelerated SARS-CoV-2 intrahost evolution leading to distinct genotypes during chronic infection. Cell Rep Med. 2023;4(2):100943. doi:10.1016/j.xcrm.2023.100943

67. Kumata R, Sasaki A. Antigenic escape is accelerated by the presence of immunocompromised hosts. Proc Biol Sci. 2022;289(1986):20221437. doi:10.1098/rspb.2022.1437

68. Harari S, Tahor M, Rutsinsky N, et al. Drivers of adaptive evolution during chronic SARS-CoV-2 infections. Nat Med. 2022;28(7):1501–1508. doi:10.1038/s41591-022-01882-4

69. El Moussaoui M, Bontems S, Meex C, et al. Intrahost evolution leading to distinct lineages in the upper and lower respiratory tracts during SARS-CoV-2 prolonged infection. Virus Evol. 2024;10(1):veae073. Published 2024 Aug 31. doi:10.1093/ve/veae073

70. Karim F, Moosa MY, Gosnell B, et al. Persistent SARS-CoV-2 infection and intra-host evolution in association with advanced HIV infection. Published online June 4, 2021. 10.1101/2021.06.03.21258228

71. Ko SH, Radecki P, Belinky F, et al. Rapid intra-host diversification and evolution of SARS-CoV-2 in advanced HIV infection. Nat Commun. 2024;15(1):7240. Published 2024 Aug 22. doi:10.1038/s41467-024-51539-8

72. Gonzalez-Reiche AS, Alshammary H, Schaefer S, et al. Sequential intrahost evolution and onward transmission of SARS-CoV-2 variants. Nat Commun. 2023;14(1):3235. Published 2023 Jun 3. doi:10.1038/s41467-023-38867-x

73. Machkovech HM, Hahn AM, Garonzik Wang J, et al. Persistent SARS-CoV-2 infection: significance and implications. Lancet Infect Dis. 2024;24(7):e453–e462. doi:10.1016/S1473-3099(23)00815-0

74. Markov PV, Ghafari M, Beer M, et al. The evolution of SARS-CoV-2. Nat Rev Microbiol. 2023;21(6):361–379. doi:10.1038/s41579-023-00878-2

75. Wilkinson SAJ, Richter A, Casey A, et al. Recurrent SARS-CoV-2 mutations in immunodeficient patients. Virus Evol. 2022;8(2):veac050. Published 2022 Aug 11. doi:10.1093/ve/veac050

76. Roemer C, Sheward DJ, Hisner R, et al. SARS-CoV-2 evolution in the Omicron era. Nat Microbiol. 2023;8(11):1952–1959. doi:10.1038/s41564-023-01504-w

77. Hill V, Du Plessis L, Peacock TP, et al. The origins and molecular evolution of SARS-CoV-2 lineage B.1.1.7 in the UK [published correction appears in Virus Evol. 2022 Dec 30;8(2):veac119. doi: 10.1093/ve/veac119.]. Virus Evol. 2022;8(2):veac080. Published 2022 Aug 26. doi:10.1093/ve/veac080

78. Qing E, Kicmal T, Kumar B, et al. Dynamics of SARS-CoV-2 Spike Proteins in Cell Entry: Control Elements in the Amino-Terminal Domains. mBio. 2021;12(4):e0159021. doi:10.1128/mBio.01590-21

79. Meng B, Datir R, Choi J, et al. SARS-CoV-2 spike N-terminal domain modulates TMPRSS2-dependent viral entry and fusogenicity. Cell Rep. 2022;40(7):111220. doi:10.1016/j.celrep.2022.111220

80. Bendall EE, Callear AP, Getz A, et al. Rapid transmission and tight bottlenecks constrain the evolution of highly transmissible SARS-CoV-2 variants. Nat Commun. 2023;14(1):272. Published 2023 Jan 17. doi:10.1038/s41467-023-36001-5

81. Sinclair P, Zhao L, Beggs CB, Illingworth CJR. The airborne transmission of viruses causes tight transmission bottlenecks. Nat Commun. 2024;15(1):3540. Published 2024 Apr 26. doi:10.1038/s41467-024-47923-z

82. Gao W, Wang L, Ju X, et al. The Deubiquitinase USP29 Promotes SARS-CoV-2 Virulence by Preventing Proteasome Degradation of ORF9b. mBio. 2022;13(3):e0130022. doi:10.1128/mbio.01300-22

83. Davis ME, Gack MU. Ubiquitination in the antiviral immune response. Virology. 2015;479–480:52-65. doi:10.1016/j.virol.2015.02.033

84. Yu M, Li J, Gao W, Li Z, Zhang W. Multiple E3 ligases act as antiviral factors against SARS-CoV-2 via inducing the ubiquitination and degradation of ORF9b. J Virol. 2024;98(6):e0162423. doi:10.1128/jvi.01624-23

85. Patchett S, Lv Z, Rut W, et al. A molecular sensor determines the ubiquitin substrate specificity of SARS-CoV-2 papain-like protease. Cell Rep. 2021;36(13):109754. doi:10.1016/j.celrep.2021.109754

86. Mols VC, Lamers MM, Leijten LM, et al. Intestinal Tropism of a Betacoronavirus (Merbecovirus) in Nathusius’s Pipistrelle Bat (Pipistrellus nathusii), Its Natural Host [published correction appears in J Virol. 2023 May 31;97(5):e0059623. doi: 10.1128/jvi.00596-23]. J Virol. 2023;97(3):e0009923. doi:10.1128/jvi.00099-23

87. Suarez R, Gregory DA, Baker DA, et al. Detecting SARS-CoV-2 cryptic lineages using publicly available whole genome wastewater sequencing data. PLoS Pathog. 2025;21(6):e1012850. Published 2025 Jun 9. doi:10.1371/journal.ppat.1012850

88. Weingarten-Gabbay S, Klaeger S, Sarkizova S, et al. Profiling SARS-CoV-2 HLA-I peptidome reveals T cell epitopes from out-of-frame ORFs. Cell. 2021;184(15):3962–3980.e17. doi:10.1016/j.cell.2021.05.046

89. Bouhaddou M, Reuschl AK, Polacco BJ, et al. SARS-CoV-2 variants evolve convergent strategies to remodel the host response. Cell. 2023;186(21):4597–4614.e26. doi:10.1016/j.cell.2023.08.026

90. Seth RB, Sun L, Ea CK, Chen ZJ. Identification and characterization of MAVS, a mitochondrial antiviral signaling protein that activates NF-kappaB and IRF 3. Cell. 2005;122(5):669–682. doi:10.1016/j.cell.2005.08.012

91. Li W, Shi Z, Yu M, et al. Bats are natural reservoirs of SARS-like coronaviruses. Science. 2005;310(5748):676–679. doi:10.1126/science.1118391

