## Supplementary Tables S1-S4 for "Genetic Evidence Indicates the Evolutionary Importance of the SARS-CoV-2 ORF9b protein"

**Supplementary Table S1**: A list of mutations used to search through large datasets for chronic infection-candidate sequences.

| Chronic Search Mutations |
|---|
| ORF1a:T12A |
| ORF1a:G30S |
| ORF1a:V38A |
| ORF1a:E55K |
| ORF1a:E57K |
| ORF1a:M85N |
| ORF1a:M85K |
| ORF1a:V86F |
| ORF1a:E93K |
| ORF1a:I95T |
| ORF1a:E102Q |
| ORF1a:E102K |
| ORF1a:E102A |
| ORF1a:E102V |
| ORF1a:I114T |
| ORF1a:L122F |
| ORF1a:H165Y |
| ORF1a:S166G |
| ORF1a:S167G |
| ORF1a:T170I |
| ORF1a:A260V |
| ORF1a:P286L |
| ORF1a:K290R |
| ORF1a:I368V |
| ORF1a:L384V |
| ORF1a:K816R |
| ORF1a:S1272N |
| ORF1a:S1272G |
| ORF1a:S1272R |
| ORF1a:D1273N |
| ORF1a:D1273G |
| ORF1a:R1318G |
| ORF1a:T1322I |
| ORF1a:T1322P |
| ORF1a:T1322A |
| ORF1a:D1323N |
| ORF1a:D1323A |
| ORF1a:D1323G |
| ORF1a:N1324K |
| ORF1a:N1324D |
| ORF1a:N1324H |
| ORF1a:N1324T |

| |
|---|
| ORF1a:N1324S |
| ORF1a:I1326V |
| ORF1a:S1361P |
| ORF1a:Q1365K |
| ORF1a:Q1365R |
| ORF1a:Q1365P |
| ORF1a:E1366G |
| ORF1a:E1366A |
| ORF1a:I1367L |
| ORF1a:I1367V |
| ORF1a:L1368I |
| ORF1a:G1369R |
| ORF1a:S1372V |
| ORF1a:S1372A |
| ORF1a:S1372P |
| ORF1a:S1372C |
| ORF1a:E1394D |
| ORF1a:T1395I |
| ORF1a:T1430P |
| ORF1a:N1458S |
| ORF1a:H1500Y |
| ORF1a:T1538I |
| ORF1a:T1542I |
| ORF1a:K1569R |
| ORF1a:V1570A |
| ORF1a:T1572K |
| ORF1a:T1638I |
| ORF1a:T1638N |
| ORF1a:T1638A |
| ORF1a:T1638P |
| ORF1a:D1639E |
| ORF1a:D1639H |
| ORF1a:D1639A |
| ORF1a:D1639N |
| ORF1a:P1640F |
| ORF1a:Y1646F |
| ORF1a:T1682A |
| ORF1a:A1708G |
| ORF1a:A1708S |
| ORF1a:N1709H |
| ORF1a:N1709T |
| ORF1a:N1709D |
| ORF1a:N1709S |
| ORF1a:F1710L |
| ORF1a:C1711S |
| ORF1a:I1714T |
| ORF1a:I1714S |
| ORF1a:I1714A |
| ORF1a:I1714V |
| ORF1a:I1714L |
| ORF1a:L1715F |

| |
|---|
| ORF1a:L1715V |
| ORF1a:A1716T |
| ORF1a:L1748V |
| ORF1a:F1779L |
| ORF1a:K1795Q |
| ORF1a:T1822I |
| ORF1a:S2024L |
| ORF1a:R2115I |
| ORF1a:D2136A |
| ORF1a:D2136G |
| ORF1a:T2154I |
| ORF1a:A2325V |
| ORF1a:A2325T |
| ORF1a:F2328V |
| ORF1a:F2328L |
| ORF1a:L2329F |
| ORF1a:L2349V |
| ORF1a:A2355G |
| ORF1a:N2370K |
| ORF1a:D2467E |
| ORF1a:E2468D |
| ORF1a:V2469A |
| ORF1a:A2470S |
| ORF1a:R2471G |
| ORF1a:R2471S |
| ORF1a:R2471K |
| ORF1a:D2472E |
| ORF1a:L2475I |
| ORF1a:D2592G |
| ORF1a:S2706G |
| ORF1a:N2708S |
| ORF1a:N2708D |
| ORF1a:A2710V |
| ORF1a:A2710G |
| ORF1a:S2926Y |
| ORF1a:S2926F |
| ORF1a:I2961F |
| ORF1a:S2972P |
| ORF1a:D2980N |
| ORF1a:D2980G |
| ORF1a:T3058I |
| ORF1a:A3070V |
| ORF1a:G3072C |
| ORF1a:H3076Y |
| ORF1a:V3077A |
| ORF1a:F3085S |
| ORF1a:R3163I |
| ORF1a:R3164H |
| ORF1a:L3201P |
| ORF1a:F3201S |
| ORF1a:D3222G |

| |
|---|
| ORF1a:D3222N |
| ORF1a:T3224A |
| ORF1a:L3249I |
| ORF1a:L3249V |
| ORF1a:I3255V |
| ORF1a:T3258N |
| ORF1a:T3287I |
| ORF1a:E3441D |
| ORF1a:N3443K |
| ORF1a:A3454V |
| ORF1a:A3456V |
| ORF1a:K3573Q |
| ORF1a:H3580Y |
| ORF1a:F3605L |
| ORF1a:L3606V |
| ORF1a:A3648T |
| ORF1a:A3648V |
| ORF1a:A3648S |
| ORF1a:A3648G |
| ORF1a:V3653A |
| ORF1a:Y3654H |
| ORF1a:R3802H |
| ORF1a:R3802C |
| ORF1a:Y3803H |
| ORF1a:T3807P |
| ORF1a:L3808F |
| ORF1a:Q3826R |
| ORF1a:Q3826K |
| ORF1a:Q3890H |
| ORF1a:V3917G |
| ORF1a:D4085E |
| ORF1a:T4087I |
| ORF1a:T4087M |
| ORF1a:T4087A |
| ORF1a:T4087V |
| ORF1a:T4088I |
| ORF1a:T4090I |
| ORF1a:E4097G |
| ORF1a:E4097V |
| ORF1a:E4097A |
| ORF1a:E4097K |
| ORF1a:E4097D |
| ORF1a:I4098V |
| ORF1a:I4098L |
| ORF1a:I4098T |
| ORF1a:Q4099H |
| ORF1a:Q4099L |
| ORF1a:Q4100H |
| ORF1a:Q4100R |
| ORF1a:V4101I |
| ORF1a:V4101F |

| |
|---|
| ORF1a:V4102I |
| ORF1a:D4117N |
| ORF1a:N4118T |
| ORF1a:P4120S |
| ORF1a:P4120L |
| ORF1a:T4164I |
| ORF1a:T4164N |
| ORF1a:D4165V |
| ORF1a:D4165G |
| ORF1a:D4165A |
| ORF1a:D4165N |
| ORF1a:D4166E |
| ORF1a:D4166S |
| ORF1a:D4166A |
| ORF1a:D4166Y |
| ORF1a:D4166G |
| ORF1a:D4166N |
| ORF1a:N4167T |
| ORF1a:N4167S |
| ORF1a:N4167K |
| ORF1a:L4169S |
| ORF1a:K4176R |
| ORF1a:P4197S |
| ORF1a:T4207A |
| ORF1a:Q4289R |
| ORF1a:T4311I |
| ORF1a:N4358K |
| ORF1a:D4395E |
| ORF1a:A4396V |
| ORF1a:S4398L |
| ORF1a:S4398P |
| ORF1a:S4398F |
| ORF1a:F4399L |
| ORF1b:E127G |
| ORF1b:V157L |
| ORF1b:V157A |
| ORF1b:L163S |
| ORF1b:S220N |
| ORF1b:Q435K |
| ORF1b:A440V |
| ORF1b:C455Y |
| ORF1b:L535I |
| ORF1b:Y610F |
| ORF1b:A679T |
| ORF1b:N725D |
| ORF1b:T730I |
| ORF1b:K774R |
| ORF1b:V783I |
| ORF1b:M785L |
| ORF1b:M785I |
| ORF1b:S786P |

| |
|---|
| ORF1b:E787A |
| ORF1b:C790Y |
| ORF1b:P821S |
| ORF1b:Y822H |
| ORF1b:D824S |
| ORF1b:D824E |
| ORF1b:D824N |
| ORF1b:D870A |
| ORF1b:D979G |
| ORF1b:V983A |
| ORF1b:Q985H |
| ORF1b:Q985R |
| ORF1b:Q985L |
| ORF1b:Q985K |
| ORF1b:L986I |
| ORF1b:Y987C |
| ORF1b:Y987H |
| ORF1b:I1118M |
| ORF1b:H1213Y |
| ORF1b:T1424I |
| ORF1b:L2019I |
| ORF1b:D2020E |
| ORF1b:I2147T |
| ORF1b:T2537I |
| ORF1b:P2648T |
| S:V3G |
| S:P9S |
| S:P9L |
| S:S13I |
| S:C15F |
| S:C15Y |
| S:C15R |
| S:T19A |
| S:T19K |
| S:T20P |
| S:P25T |
| S:P25L |
| S:F32S |
| S:K41E |
| S:V42I |
| S:V42F |
| S:L48S |
| S:L48V |
| S:W64R |
| S:W64G |
| S:W64L |
| S:W64C |
| S:H66Q |
| S:H69N |
| S:V70T |
| S:N74T |

| |
|---|
| S:D80G |
| S:I101T |
| S:E132Q |
| S:C136R |
| S:C136F |
| S:N137M |
| S:P139S |
| S:P139T |
| S:G142C |
| S:Y144C |
| S:Y144D |
| S:H146D |
| S:S151N |
| S:S151C |
| S:S151G |
| S:W152C |
| S:W152L |
| S:E154K |
| S:R158S |
| S:R158I |
| S:K187E |
| S:K187T |
| S:R190K |
| S:N196S |
| S:D198N |
| S:F201L |
| S:K206N |
| S:I210T |
| S:G213E |
| S:R214S |
| S:R214C |
| S:R214H |
| S:D215G |
| S:P230S |
| S:I231T |
| S:G232D |
| S:H245Y |
| S:Y248F |
| S:G257D |
| S:T274I |
| S:T274N |
| S:T299I |
| S:V327A |
| S:P330S |
| S:N334K |
| S:L335S |
| S:P337N |
| S:P337S |
| S:P337R |
| S:D339A |
| S:D339H |

| |
|---|
| S:D339Y |
| S:H339R |
| S:D339V |
| S:D339E |
| S:E340K |
| S:E340A |
| S:E340V |
| S:E340Q |
| S:E340D |
| S:V341I |
| S:N343S |
| S:A344V |
| S:T345I |
| S:T345A |
| S:R346K |
| S:R346I |
| S:K346E |
| S:R346S |
| S:K346T |
| S:A348P |
| S:N354K |
| S:N354D |
| S:K356E |
| S:R357T |
| S:R357K |
| S:I358F |
| S:N360H |
| S:D364N |
| S:V367A |
| S:V367F |
| S:Y369H |
| S:Y369D |
| S:Y369N |
| S:N370K |
| S:L371F |
| S:A372S |
| S:A372T |
| S:P373T |
| S:F374L |
| S:F374S |
| S:F375Y |
| S:A376V |
| S:T376N |
| S:A376P |
| S:F377L |
| S:K378R |
| S:K378T |
| S:K378E |
| S:K378N |
| S:P384L |
| S:P384S |

| |
|---|
| S:P384A |
| S:T385I |
| S:N388D |
| S:V395I |
| S:R403S |
| S:N405S |
| S:E406Q |
| S:T415A |
| S:N417H |
| S:N417D |
| S:N417I |
| S:K417T |
| S:N417Y |
| S:N417T |
| S:D420N |
| S:I434M |
| S:N440D |
| S:K440R |
| S:K444E |
| S:K444S |
| S:K444N |
| S:P445T |
| S:H445R |
| S:V445A |
| S:P445L |
| S:P445H |
| S:V445L |
| S:P445S |
| S:V445S |
| S:S446R |
| S:G446R |
| S:G446N |
| S:G446V |
| S:G446D |
| S:N448S |
| S:Y449H |
| S:Y449N |
| S:Y449S |
| S:N450K |
| S:W452R |
| S:R452Q |
| S:L452Q |
| S:Q452K |
| S:Y453F |
| S:L455W |
| S:S455W |
| S:S455A |
| S:F456V |
| S:N460T |
| S:L461I |
| S:K462E |

| |
|---|
| S:K462T |
| S:T470N |
| S:Y473F |
| S:G476S |
| S:N477D |
| S:K478R |
| S:K478E |
| S:G482S |
| S:V483A |
| S:A484K |
| S:A484V |
| S:E484Q |
| S:A484P |
| S:E484D |
| S:E484V |
| S:E484R |
| S:A484T |
| S:G485S |
| S:F486I |
| S:V486A |
| S:V486P |
| S:F486L |
| S:P486H |
| S:S490Y |
| S:F490Y |
| S:F490V |
| S:F490L |
| S:R493G |
| S:Q493K |
| S:Q493L |
| S:Q493V |
| S:S494P |
| S:S496T |
| S:G496D |
| S:G496N |
| S:G496V |
| S:S496N |
| S:Q498H |
| S:Q498K |
| S:Q498Y |
| S:P499R |
| S:T500G |
| S:T500S |
| S:T500N |
| S:T500A |
| S:Y501F |
| S:N501T |
| S:G504S |
| S:G504D |
| S:H505N |
| S:Y508H |

| |
|---|
| S:H519R |
| S:H519Q |
| S:H519N |
| S:A522P |
| S:K529N |
| S:V551I |
| S:Q564E |
| S:F565S |
| S:G566S |
| S:R567S |
| S:I569M |
| S:A570T |
| S:D571G |
| S:T572N |
| S:T573I |
| S:D574N |
| S:D574V |
| S:L582F |
| S:F592S |
| S:G593V |
| S:L611F |
| S:Q613R |
| S:P621A |
| S:P621T |
| S:I624M |
| S:H625N |
| S:H625R |
| S:Q628K |
| S:L629V |
| S:R634H |
| S:V642G |
| S:R646S |
| S:A647V |
| S:A653V |
| S:N657H |
| S:N657K |
| S:S659P |
| S:S659A |
| S:H681R |
| S:R683Q |
| S:T732I |
| S:S735L |
| S:S735A |
| S:D737E |
| S:D737Y |
| S:M740I |
| S:S758G |
| S:Q762E |
| S:N764S |
| S:R765L |
| S:G769E |

| |
|---|
| S:T791I |
| S:K795T |
| S:P812R |
| S:L828F |
| S:Q836K |
| S:G838S |
| S:D839G |
| S:D839N |
| S:D839A |
| S:D839H |
| S:D843G |
| S:A852V |
| S:A852K |
| S:K854R |
| S:K854E |
| S:K854N |
| S:T859N |
| S:G885F |
| S:G885T |
| S:I896T |
| S:D936H |
| S:D936Y |
| S:D936G |
| S:T941K |
| S:A942E |
| S:A942T |
| S:A944V |
| S:A944T |
| S:A944S |
| S:Q949L |
| S:Q949R |
| S:V952F |
| S:A958S |
| S:A958D |
| S:N960D |
| S:N960S |
| S:N960T |
| S:V963F |
| S:S968A |
| S:K969T |
| S:V976A |
| S:N978D |
| S:N978T |
| S:N978S |
| S:D979E |
| S:L981V |
| S:S982L |
| S:D985N |
| S:V987F |
| S:Q1002H |
| S:S1003I |

| |
|---|
| S:T1027I |
| S:H1058Y |
| S:H1101D |
| S:H1101R |
| S:P1143S |
| S:I1169T |
| S:A1174T |
| S:S1175P |
| S:I1179T |
| S:I1179S |
| S:E1182G |
| S:I1183T |
| S:I1183F |
| S:R1185S |
| S:L1186F |
| S:V1189A |
| S:E1258Q |
| S:E1258A |
| E:V5L |
| E:V5A |
| E:V5F |
| E:S6L |
| E:G10S |
| E:G10C |
| E:G10V |
| E:N15S |
| E:S16N |
| E:L18R |
| E:L18I |
| E:L19F |
| E:F20S |
| E:F23S |
| E:F23V |
| E:F23L |
| E:L27S |
| E:L28P |
| E:T30I |
| E:L31I |
| E:A32V |
| E:T35I |
| E:A36V |
| E:L37H |
| E:Y42C |
| E:Y42F |
| E:C44R |
| E:N48S |
| E:S50I |
| E:S55F |
| E:V58A |
| E:V62I |
| M:A2V |

| |
|---|
| M:D3Y |
| M:H3L |
| M:N3S |
| M:N3K |
| M:D3A |
| M:N3I |
| M:S4F |
| M:S4L |
| M:S4P |
| M:G6C |
| M:G6S |
| M:T7G |
| M:I8F |
| M:I8S |
| M:I8T |
| M:I8L |
| M:E11V |
| M:E12A |
| M:E12G |
| M:L16F |
| M:L17V |
| M:L17F |
| M:E18D |
| M:Q19H |
| M:I24L |
| M:F28S |
| M:A40P |
| M:I48M |
| M:A69V |
| M:I73V |
| M:T77N |
| M:S94R |
| M:S94N |
| M:S99A |
| M:R107S |
| M:H125Y |
| M:L138I |
| M:I144V |
| M:H148R |
| M:H155Q |
| M:H155N |
| M:S173P |
| M:A188T |
| M:G189S |
| M:S197T |
| N:D3L |
| N:L13F |
| N:S37P |
| N:A134V |
| N:T148A |
| N:I157V |

| |
|---|
| N:T271I |
| N:P326H |
| N:P326F |
| N:P326L |
| N:S327L |
| ORF3a:I10L |
| ORF3a:H182D |
| ORF3a:Q213K |
| ORF3a:Q213R |
| ORF3a:P274S |
| ORF6:P57S |
| ORF7a:E22D |
| ORF7a:R39I |
| ORF7a:T39I |
| ORF7a:N43S |
| ORF7a:N43K |
| ORF7a:F59I |
| ORF7a:S60T |
| ORF7a:T61S |
| ORF7a:R78H |
| ORF7a:S81P |
| ORF7a:L96P |
| ORF7a:L96R |
| ORF7a:A105V |
| ORF7a:T115I |
| ORF8:V117L |
| ORF9b:F10S |
| ORF9b:A11E |
| ORF9b:L12F |
| ORF9b:R13H |
| ORF9b:R13C |
| ORF9b:Q18* |
| ORF9b:L64P |
| ORF9b:L64R |
| ORF9b:Q77E |
| ORF9b:K80E |
| ORF9b:E86D |
| ORF9b:D89G |
| ORF9b:D89E |
| ORF9b:E90G |
| ORF9b:V92M |
| ORF9b:V93L |
| ORF9b:V93M |
| ORF9b:T95M |
| ORF9b:T95A |
| ORF9b:V96A |
| ORF9b:V96E |
| ORF9b:K97E |

**Supplementary Table S2**: Amino acid substitution density for each canonical protein in EPCI and HQCS datasets. The ratio of these densities, after being adjusted for the greater total number of

| Gene | AA Substitution Density EPCI | AA Substitution Density HCQS | Chronic-to-Circulating AA Substitution Ratio |
|---|---|---|---|
| NSP1 | 7.59 | 2.5 | 9.43 |
| NSP2 | 3.37 | 2.5 | 4.19 |
| NSP3 | 4.52 | 1.8 | 7.80 |
| NSP4 | 3.66 | 1.3 | 8.74 |
| NSP5 | 2.36 | 1.1 | 6.66 |
| NSP6 | 4.79 | 1.8 | 8.26 |
| NSP7 | 2.92 | 1.0 | 9.07 |
| NSP8 | 5.64 | 1.2 | 14.59 |
| NSP9 | 8.25 | 1.5 | 17.08 |
| NSP10 | 2.63 | 1.1 | 7.42 |
| NSP12 | 2.17 | 1.0 | 6.74 |
| NSP13 | 2.62 | 1.2 | 6.78 |
| NSP14 | 1.76 | 1.3 | 4.20 |
| NSP15 | 1.97 | 1.5 | 4.08 |
| NSP16 | 1.94 | 1.3 | 4.63 |
| ORF3a | 6.84 | 5.1 | 4.16 |
| ORF6 | 4.52 | 2.0 | 7.02 |
| ORF7a | 9.99 | 4.6 | 6.74 |
| ORF7b | 4.93 | 2.9 | 5.28 |
| ORF8 | 8.05 | 5.2 | 4.81 |
| ORF9b | 16.7 | 4.0 | 12.96 |
| S | 14.09 | 1.8 | 24.30 |
| E | 15.57 | 1.2 | 40.28 |
| M | 7.25 | 1.2 | 18.76 |
| N | 6.39 | 2.7 | 7.35 |

**Supplementary Table S3**: Most enriched ORF9b amino acid substitutions in EPCI dataset relative to HQCS dataset (EPCI sequence count ≥ 5). Due to the enormous number of artifactual reversions in sequence databases, reversion substitutions were not counted in the HQCS dataset. They are placed at the top because the authors judge they would be at or near the top if it was possible to ascertain the number of genuine reversions in HQCS sequences.

| Rank | Substitution | EPCI:HQCS private AA substitution ratio adjusted for sequence & mutation count | Number of Independent Substitutions | |
|---|---|---|---|---|
| | | | EPCI | HQCS |
| 1 | ORF9b:G16D | NA (reversion) | 54 | NA |
| 2 | ORF9b:S10P | NA (reversion) | 50 | NA |
| 3 | ORF9b:K97E | 519.60 | 71 | 94 |
| 4 | ORF9b:A11E | 152.87 | 12 | 54 |
| 5 | ORF9b:V93L | 118.20 | 111 | 646 |
| 6 | ORF9b:F10S | 103.19 | 9 | 60 |
| 7 | ORF9b:D89E | 63.81 | 100 | 1078 |
| 8 | ORF9b:T95A | 56.96 | 26 | 314 |
| 9 | ORF9b:V96A | 42.77 | 24 | 386 |
| 10 | ORF9b:V93M | 38.55 | 26 | 464 |
| 11 | ORF9b:V96E | 34.98 | 6 | 118 |
| 12 | ORF9b:K80E | 32.98 | 7 | 146 |
| 13 | ORF9b:R13H | 24.16 | 38 | 1082 |
| 14 | ORF9b:Q77E | 17.83 | 7 | 270 |
| 15 | ORF9b:L64P | 17.25 | 73 | 2912 |
| 16 | ORF9b:L64R | 16.88 | 8 | 326 |
| 17 | ORF9b:V92M | 16.59 | 11 | 456 |
| 18 | ORF9b:R13C | 9.55 | 42 | 3026 |
| 19 | ORF9b:L12F | 9.44 | 11 | 802 |
| 20 | ORF9b:V94L | 9.21 | 9 | 672 |
| 21 | ORF9b:Q34* | 8.29 | 8 | 664 |
| 22 | ORF9b:K67E | 7.97 | 7 | 604 |
| 23 | ORF9b:K67R | 7.93 | 11 | 954 |
| 24 | ORF9b:T95M | 7.38 | 72 | 6710 |
| 25 | ORF9b:P10S | 7.28 | 38 | 3592 |
| 26 | ORF9b:R25S | 6.29 | 6 | 656 |
| 27 | ORF9b:M8V | 6.13 | 7 | 786 |
| 28 | ORF9b:Q77* | 6.08 | 10 | 1132 |
| 29 | ORF9b:D66G | 6.06 | 5 | 568 |
| 30 | ORF9b:N62D | 5.98 | 8 | 920 |

| | | | | |
|---|---|---|---|---|
| 31 | ORF9b:A68E | 5.76 | 6 | 716 |
| 32 | ORF9b:L61S | 5.53 | 8 | 996 |
| 33 | ORF9b:M1T | 5.33 | 7 | 904 |
| 34 | ORF9b:L71S | 4.76 | 5 | 722 |
| 35 | ORF9b:M1I | 4.46 | 7 | 1080 |
| 36 | ORF9b:Q18H | 4.44 | 14 | 2170 |
| 37 | ORF9b:M1V | 4.28 | 6 | 964 |
| 38 | ORF9b:E65Q | 4.20 | 5 | 818 |
| 39 | ORF9b:I5T | 4.11 | 70 | 11710 |
| 40 | ORF9b:P17S | 3.96 | 16 | 2776 |
| 41 | ORF9b:E86D | 3.86 | 77 | 13716 |
| 42 | ORF9b:P51Q | 3.64 | 8 | 1512 |
| 43 | ORF9b:T84I | 3.59 | 6 | 1150 |
| 44 | ORF9b:M8T | 3.47 | 5 | 992 |
| 45 | ORF9b:A68V | 2.37 | 10 | 2902 |
| 46 | ORF9b:M78T | 2.28 | 12 | 3618 |
| 47 | ORF9b:S6N | 2.20 | 10 | 3126 |
| 48 | ORF9b:L64F | 2.17 | 23 | 7296 |
| 49 | ORF9b:A11T | 2.08 | 6 | 1984 |
| 50 | ORF9b:D2E | 2.06 | 14 | 4664 |

**Supplementary Table S4**: Sequences with two or more private mutations in the last nine residues of ORF9b:89-97. EPCI = Expanded Posited Chronic-Infection dataset, HQCS = High-Quality Circulating Sequences (see Methods),

HQCS relaxed QC = Nextclade qc score ≤30, private AA substitutions ≤10, private reversions ≤ 1, <150 total nucleotide deletions

| Dataset | ≥2 ORF9b:89-97 AA Substitutions | <2 ORF9b:89-97 private AA Substitutions | ≥2 ORF9b:89-97 AA Substitutions % | EPCI Fold-Enrichment |
|---|---|---|---|---|
| EPCI | 68 | 3150 | 2.11% | ref |
| HQCS | 9 | 9990952 | 0.0000901% | 23964 |
| HQCS relaxed QC | 26 | 13833331 | 0.000188% | 11486 |
